# Ion Mobility-Enhanced LA-REIMS Improves Molecular Resolution in Ambient Biofluid Metabolomics

**DOI:** 10.64898/2026.03.10.709786

**Authors:** Vera Plekhova, Noa Van de Velde, Anne VandenBerghe, José Diana Di Mavungu, Lynn Vanhaecke

## Abstract

Ambient metabolomics techniques such as laser-assisted rapid evaporative ionization mass spectrometry (LA-REIMS) enable fast, preparation-free fingerprinting of biological samples but are inherently limited by spectral congestion in the absence of chromatographic separation. While ion mobility spectrometry provides additional gas-phase separation, maintaining ion transmission under the transient signals’ characteristic of laser desorption, remains analytically challenging. Here, we define operating conditions for cyclic traveling-wave ion mobility spectrometry (cIMS) that preserve transmission under LA-REIMS duty-cycle constraints and systematically evaluate how cIMS integration reshapes biofluid fingerprints and enhances chemical specificity in chromatography-free metabolomics analysis.

Under optimized single-pass conditions, cIMS separation reorganized LA-REIMS spectra into structured mass/mobility feature domains, enabling selective mobility-based filtering of matrix-derived salt cluster ions. This reduced non-biological background contributions by up to 35% of total spectral intensity while preserving over 90% of detected untargeted features. Although cIMS operation introduced a sensitivity penalty relative to time-of-flight-only acquisition, approximately 80% of the total ion current was recovered under optimized conditions. Mobility-resolved data revealed coherent homologous series and class-specific structural trends, particularly for lipids, supporting class-level annotation. Analysis of 101 metabolite and lipid standards covering a broad physicochemical range (logP −5.30 to 19.40) demonstrated comprehensive molecular coverage, high mass accuracy (mean 2.4 ppm), and good agreement with reference CCS values (mean deviation 4.0%), with isomer separation observed for biologically important secondary bile acids in extended separation cycles. Collectively, these results establish LA-REIMS-cIMS as a practical analytical strategy for enhancing chemical specificity and spectral interpretability in support of high-throughput large-scale metabolic fingerprinting.

Graphical abstract
Ion mobility spectrometry adds an orthogonal gas-phase separation to LA-REIMS, reorganizing complex biofluid spectra into distinct mass-mobility feature bands and improving molecular resolution in rapid ambient ionization metabolomics.

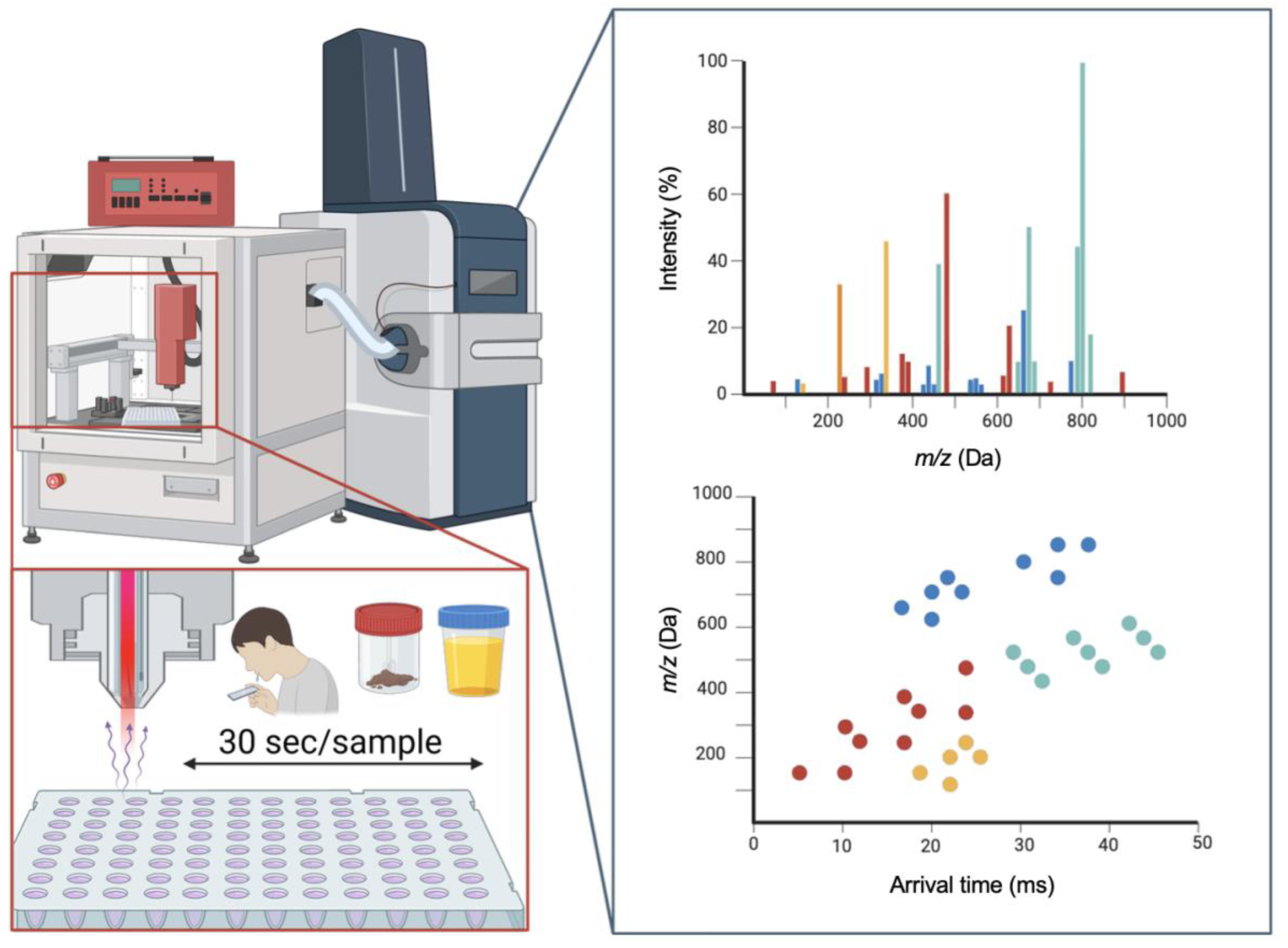

## Introduction

Ambient ionization mass spectrometry (AIMS) enables rapid molecular fingerprinting of biological samples with minimal or no sample preparation and without chromatographic separation, in contrast to conventional metabolomics workflows^1,2^. These characteristics support high analytical throughput and operational simplicity, making AIMS particularly attractive for in-field applications, large-sample-number metabolomics studies, and workflows requiring fast, on-site analytical feedback, such as epidemiological screening, environmental monitoring, and manufacturing process control, among others^1,3,4^. In biomedical contexts specifically, AIMS has attracted growing interest for the sensitive molecular characterization of clinical specimens, where rapid and robust analysis can support point-of-care decision making and population-level surveillance initiatives aligned with emerging 5P medicine frameworks^5,6^.

Among AIMS techniques, laser-assisted rapid evaporative ionization mass spectrometry (LA-REIMS) occupies a distinct position due to its applicability to a wide range of biological samples, including non-invasively collected urine, saliva, and feces ^7^. LA-REIMS generates ions via rapid laser desorption and post-ionization, producing short-lived ion populations that closely reflect the bulk biochemical composition of the sample^8^. While this mechanism enables exceptional analytical speed and robustness to sample morphology, it also leads to highly congested mass spectra characterized by isobaric overlap, multiple adduct formation, and matrix-derived background signals, particularly in complex biofluids^9^. In the absence of chromatographic separation and prolonged ion accumulation, opportunities to disentangle endogenous metabolites from co-ionized matrix components are inherently limited. Improving chemical specificity without compromising throughput therefore remains a central challenge for biological fingerprint interpretation and confident compound assignment in AIMS-based biofluid metabolomics^10^.

In this context, ion mobility spectrometry (IMS) offers a promising, non-chromatographic means of increasing chemical specificity while preserving the rapid analysis characteristic of ambient ionization workflows^11^. By introducing a gas-phase separation based on ion size, shape, and charge, IMS adds an orthogonal conformational dimension to datasets acquired without prior chromatographic separation, that redistributes ions across arrival time, reducing spectral congestion and enabling more reliable association of signals with individual compounds^12^. In addition, IMS allows for the determination of collision cross section (CCS) values, which are more transferable across platforms than chromatographic retention times and can support compound annotation through standardized reference databases^13^. These attributes have established IMS as a valuable complement to mass spectrometry and motivate its systematic evaluation in AIMS workflows under the stringent duty-cycle constraints imposed by ambient ionization.

Among available IMS methods, cyclic traveling-wave ion mobility spectrometry (cTWIMS, cIMS) offers a combination of features that are particularly relevant for addressing these constraints. In uniform-field drift tube IMS, resolving power is fixed and linked to drift time (and thus drift length and operating conditions), creating an inherent limitation to resolution and separation time tuning^14^. By contrast, cTWIMS enables users to select the effective separation length by varying the number of passes around the cyclic device, increasing resolving power with longer cycle times^15^. Unlike differential mobility methods such as high-field asymmetric waveform IMS, which function as differential ion filters with transmission efficiencies that can vary strongly between ion classes, cTWIMS operates as a time-dispersive separation that can be acquired in a broadband manner^16,17^. This enables short, single-pass operation for high-throughput measurements while retaining the option to extend separations for targeted analyses when additional mobility resolution is required.

Building on these considerations, this work presents the integration of cIMS with LA-REIMS for high-throughput biofluid analysis. While cIMS has previously been applied in chromatography-free settings^12^, its implementation has not previously been adapted for ion sources generating transient ion packets, such as LA-REIMS. Here, we explore the potential of cIMS to operate in a manner compatible with transient ion generation and rapid acquisition, while providing mobility-resolved information that enhances spectral interpretability. By evaluating mobility performance, ion transmission, and duty-cycle trade-offs, and by assessing the extent to which mobility separation mitigates matrix-driven spectral complexity while enabling metabolite class annotation and isomer discrimination in the absence of chromatography, this study defines both the potential and practical limits of cIMS for fit-for-purpose integration into ambient biofluid metabolomics.

## Materials and methods

### 1. Chemical standards and solvents

Isopropyl alcohol (IPA, LC-MS grade) was purchased from Fisher Scientific (UK). Ultrapure water (UPW) was produced using a Sartorius Arium 661 UV water purification system (Sartorius, Belgium). Analytical standards were purchased from Sigma-Aldrich (USA), ICN Biomedicals Inc. (USA), TLC Pharmchem (Canada), or Waters Corporation (USA) as listed previously^18^. Sodium chloride (NaCl, food-grade purity) was dissolved in ultrapure water to yield a 0.9% (w/v) solution corresponding to isotonic saline conditions.

### 2. Samples and study design

Biological samples were obtained from existing, ethically approved clinical cohorts of which re-use was allowed via biobank BC-184. Saliva samples originated from the FAME study^19^ and urine and feces from the MetaBEAse cohort^5^. Study specimens were composed as pooled samples comprising equal parts from three normal-weight and three overweight children, classified according to International Obesity Task Force (IOTF) criteria^20^, extending chemical diversity of the biological pool by including individuals with different phenotypical traits. Pooling was applied to minimize inter-individual biological variability and to enable controlled assessment of analytical performance.

Following at-home collection, samples were immediately frozen at −18 °C. Upon receipt at LIMET (within 48 h), fecal samples were freeze-dried, homogenized by grinding and sieving, and subsequently stored at −80 °C to rapidly quench ongoing microbial, enzymatic, and chemical degradation^21^. Saliva and urine samples were stored at −80 °C without additional processing. Prior to analysis, samples were thawed at room temperature (22 ± 2 °C). Urine and saliva samples were vortex-mixed and aliquoted (100 µL) into 96-well plates for direct analysis without further purification^7^. Freeze-dried fecal samples were resuspended with UPW (1:4, w/v) to generate fecal water, facilitating homogenization and subsequent infrared laser desorption^5^. An aqueous sodium chloride solution (0.9% (w/v), physiological saline concentration) prepared in ultrapure water was analyzed in triplicate under identical LA-REIMS-cIMS conditions and used as a reference for the identification of salt-derived cluster ions in the ion mobility space.

### 3. Instrumentation and acquisition modes

LA-REIMS experiments were performed using a mid-infrared laser ablation system (Opolette™ 2940, OPOTEK, USA) coupled to a REIMS ionization source (Waters Corporation, Manchester, UK) as described previously^7^. A continuous flow (0.2 mL/min) of IPA was supplied to the source throughout all experiments to support post-ionization and ion transfer. The REIMS ionization source was coupled to a SELECT SERIES™ Cyclic™ traveling-wave ion mobility (cIMS) time-of-flight (ToF) MS (Waters Corporation, Manchester, UK)^15^. Nitrogen was used as the ion mobility drift gas. Data were acquired in both positive and negative ionization modes for targeted experiments, with polarity-specific lock-mass correction applied as described below. Untargeted biofluid fingerprinting experiments were performed in negative ionization mode only, because positive mode produced high levels of background-derived peaks.

General ion optics parameters, solvent delivery conditions, and ToF scan rates were based on previously optimized biofluid-specific LA-REIMS protocols^7^ and adapted to the instrument configuration used in this study. Detailed parameter settings are provided in SI Note 1.

### 4. cIMS operation and optimization

cIMS method optimization was performed using pooled urine as a representative biological matrix, selected due to its high molecular fingerprint reproducibility, as previously demonstrated for ToF-based LA-REIMS analyses^7^. Urine was used consistently throughout cIMS optimization to minimize matrix-related variability and enable robust comparison of experimental conditions. The cIMS mode was initially operated using default (manufacturer-set) parameters (total cycle time 51.6 ms, traveling wave (TW) velocity 375 m/s, TW height 26 V). Optimized cIMS experiments were performed using a TW voltage ramp (total cycle time 37.8 ms, TW velocity 375 m/s, TW starting height 10 V, ending height 26 V, ramping rate 1 V/ms) in combination with optimized ToF parameters (SI Note 1). The cIMS cycle time, and TW ramp starting and ending voltages were optimized iteratively during acquisition by monitoring the total ion current (TIC). TW ramping rate and wave velocity were further fine-tuned using Design of Experiments (DoE) modelling to promote a broad ion arrival-time distribution, thereby enhancing ion mobility separation and resolving power under short cycle-time conditions as detailed in SI Note 2. Benchmarking experiments were conducted using serial measurements (n = 10) of pooled urine sample aliquots under each acquisition mode to allow direct comparison of signal intensity, feature detection, and reproducibility.

### 5. Untargeted LA-REIMS(-cIMS)

Untargeted metabolomics analyses were performed on pooled urine, saliva, and fecal water samples (n = 10 per matrix). Data were acquired using both ToF-only acquisition (cIMS bypassed), default (manufacturer-set) single-pass cIMS mode and optimized single-pass cIMS conditions. Physiological saline samples were analyzed to support identification of salt-derived cluster ion bands in mobility-resolved spectra. Mobility-domain filtering was applied by excluding arrival-time regions corresponding to these clusters. Average feature counts and summed feature intensities were compared before and after filtering to assess the contribution of salt-derived clusters to the overall signal structure.

### 6. Targeted LA-REIMS-(cIMS)

A panel of 101 in-house metabolite and lipid standards spanning a wide range of molecular weights (88.02 to 1152.71 Da) and lipophilicity (logP −5.30 to 19.40), covering the major metabolome and lipidome (9 HMDB superclasses and 39 HMDB classes^22^) was analyzed to assess compound detectability, ion transmission, and annotation performance (SI Table 4). To isolate ToF and cIMS transmission effects from laser-sample interactions, standards were introduced by direct infusion into the REIMS source.

Standards were grouped into two polarity-specific mixtures (polar metabolites and lipids prepared separately based on stock solution compatibility), with each compound present at a final concentration of 10 ng/µL. When required, mixtures were reanalyzed at 100 ng/µL per compound (SI Table 4). IPA blanks were measured before analytical standard mixtures in triplicate for downstream background subtraction. Each mixture was analyzed in triplicate in both positive and negative ionization modes and detectability metrics were based on averaged measurements. Lock-mass correction was applied using cholic acid (negative mode) and anandamide C20:4 (positive mode), selected as polarity-specific references due to their consistent high-intensity signals under REIMS conditions. For each measurement, 50 µL of the analytical standard mixture was infused and the source was flushed with 500 µL IPA after each acquisition.

In ToF-only mode, compound matching was based on averaged *m/z* values using a tolerance of 10 ppm. In cIMS-enabled mode, compound matching was based on combined averaged *m/z* and collision cross section (CCS) values. Averaged *m/z* and CCS values were calculated from replicate measurements (n = 3), except for compounds used for CCS calibration, for which two replicates were averaged and one reserved exclusively for calibration. CCS calibration was performed post-acquisition with subset of standards with available literature CCS values spanning a broad *m/z* and mobility range (SI Table 3). For cIMS-based evaluation, a CCS deviation cut-off of 10% was applied to compounds with available empirical reference CCS values from open databases, as detailed in SI Table 5.

Analytical standards of secondary bile acids (deoxycholic acid, ursodeoxycholic acid, chenodeoxycholic acid and their glycine- and taurine-conjugated forms; Table 1) and positional isomers of phosphocholine lipids (PC 16:0/18:0 and PC 18:0/16:0). were analyzed at 10 ng/µL concentration in triplicate, accompanied by solvent blanks, as described above. Compounds were first analyzed individually (10 ng/µL) to record arrival times and subsequently as defined mixtures under single-pass and extended cIMS conditions (2 - 200 ms effective separation time). All other source, ion optics, and ToF parameters were held constant. Arrival-time distributions of deprotonated monomers and selected adduct or cluster ions were compared across individual and mixed analyses.

**Table 1.** Bile acid analytical standards and their detected forms with arrival times following LA-REIMS-cIMS analysis.

| Compound | Chemical Formula | Arrival time (ms) |  |  |  |
| --- | --- | --- | --- | --- | --- |
| | | $[M-H]^-$ | $[2M-H]^-$ | $[3M-H]^-$ | $[2M+Na-2H]^-$ |
| Deoxycholic acid | C24H40O4 | 31.31 | 70.47 | 22.01 | ND |
| Ursodeoxycholic acid | C24H40O4 | 32.66 | 71.24 | 22.56 | ND |
| Chenodeoxycholic acid | C24H40O4 | 32.92 | 65.39 | 21.65 | ND |
| Glycoursodeoxycholic acid | C26H43NO5 | 31.18 | 82.21 | OOD | 83.52 |
| Glycodeoxycholic acid | C26H42NO5 | 31.26 | 83.52 | OOD | 84.11 |
| Glycochenodeoxycholic acid | C26H42NO5 | 31.07 | 79.86 | OOD | 83.52 |
| Taurochenodeoxycholic acid | C26H44NO6S | 33.88 | 89.69 | OOD | 91.40 |
| Taurodeoxycholic acid | C26H44NO6S | 34.19 | 88.34 | OOD | 94.53 |
| Tauroursodeoxycholic | C26H45NO6S | 34.14 | 88.30 | OOD | 94.55 |

ND: not detected; OOD: outside of tested $m/z$ range (50 – 1200 Da)

### 7. Data processing and statistical analysis

Raw mass spectra were acquired using MassLynx (v. 4.2, Waters Corporation). Spectra containing mobility information were processed in DriftScope (v. 3.0, Waters Corporation) for individual compound detection and with MS-DIAL^23^ (v. 5.0, RIKEN) for batch processing. Progenesis Bridge and QI (v. 2.3, Waters Corporation) were used for signal alignment, background subtraction, peak picking and TIC normalization. DoE modelling^24^ for ToF and IMS parameters optimization was performed in JMP® (v. 18, SAS Institute Inc., Cary, NC, USA), fitting standard least-squares regression to assess response effects. Model effects were evaluated by analysis of variance (ANOVA), and factor significance was determined using F-tests with a significance threshold of α = 0.05 and using FDR correction. The JMP Prediction Profiler was used to visualize factor effects and to identify parameter settings that maximized feature count and repeatability based on model response predictions.

For selected untargeted analyses, mobility-domain filtering was applied by excluding arrival-time regions corresponding to salt-derived cluster ion bands identified in IMS *m/z* vs mobility arrival time maps. Feature counts and summed feature intensities were calculated before and after filtering to assess the contribution of these clusters to overall signal structure.

For targeted data analysis, .raw directories were converted to .mzML format using msConvert^25^. The resulting files were then visually curated in a custom R Shiny application to confirm the presence of the expected compound peaks (SI Note 3). Target lists were generated based on the known compound identity and expected adducts per polarity. For each replicate, 10 scans around the TIC apex were selected and summarized. Detected peaks were manually curated based on signal presence in samples relative to blanks, and detection quality metrics (mass deviation before and after lock mass correction, signal-to-noise (S/N) ratio were recorded. Curated results were exported for downstream analysis.

CCS calibration was performed using a subset of experimentally detected standards with visually confirmed peaks and reference CCS values obtained from the Ion Mobility Collision Cross Section Compendium^26^. Calibrants were selected to span a broad *m/z* and CCS range and to avoid isomeric or closely related compounds. A peak intensity threshold of 80 counts (negative mode) or 100 counts (positive mode), based on polarity-specific noise levels, and an *m/z* tolerance of 0.1 Da were applied during calibration.

## Results and discussion

### 1. Duty-cycle-aware optimization enables compatible integration of cIMS with LA-REIMS

Initial experiments were designed to evaluate the impact of cIMS incorporation on the quality of untargeted LA-REIMS biofluid fingerprints, using pooled urine (10 replicates) as a representative matrix. When cIMS was enabled using manufacturer-set default parameters, a pronounced reduction in overall sample signal was observed compared to ToF-only acquisition. The average total ion current (TIC) decreased by approximately one order of magnitude, from 2.87 × 10⁷ ± 5.24 × 10⁵ in ToF mode to 1.54 × 10⁶ ± 2.59 × 10⁵ in default cIMS mode (Fig. 1A). In addition to reduced TIC, relative peak abundances were markedly altered across the full detected mass range (50-1200 Da), with both decreases and increases in individual feature intensities observed upon cIMS activation compared to ToF-only acquisition (Fig. 1, A-C, right panel). These observations are consistent with duty-cycle and ion transmission losses associated with ion mobility operation under the transient ion generation conditions characteristic of LA-REIMS, indicating that default cIMS settings optimized for more continuous ion sources are not directly compatible with ambient ionization workflows^27^. This prompted systematic optimization toward duty-cycle-aware cIMS operation.

**Fig. 1.**
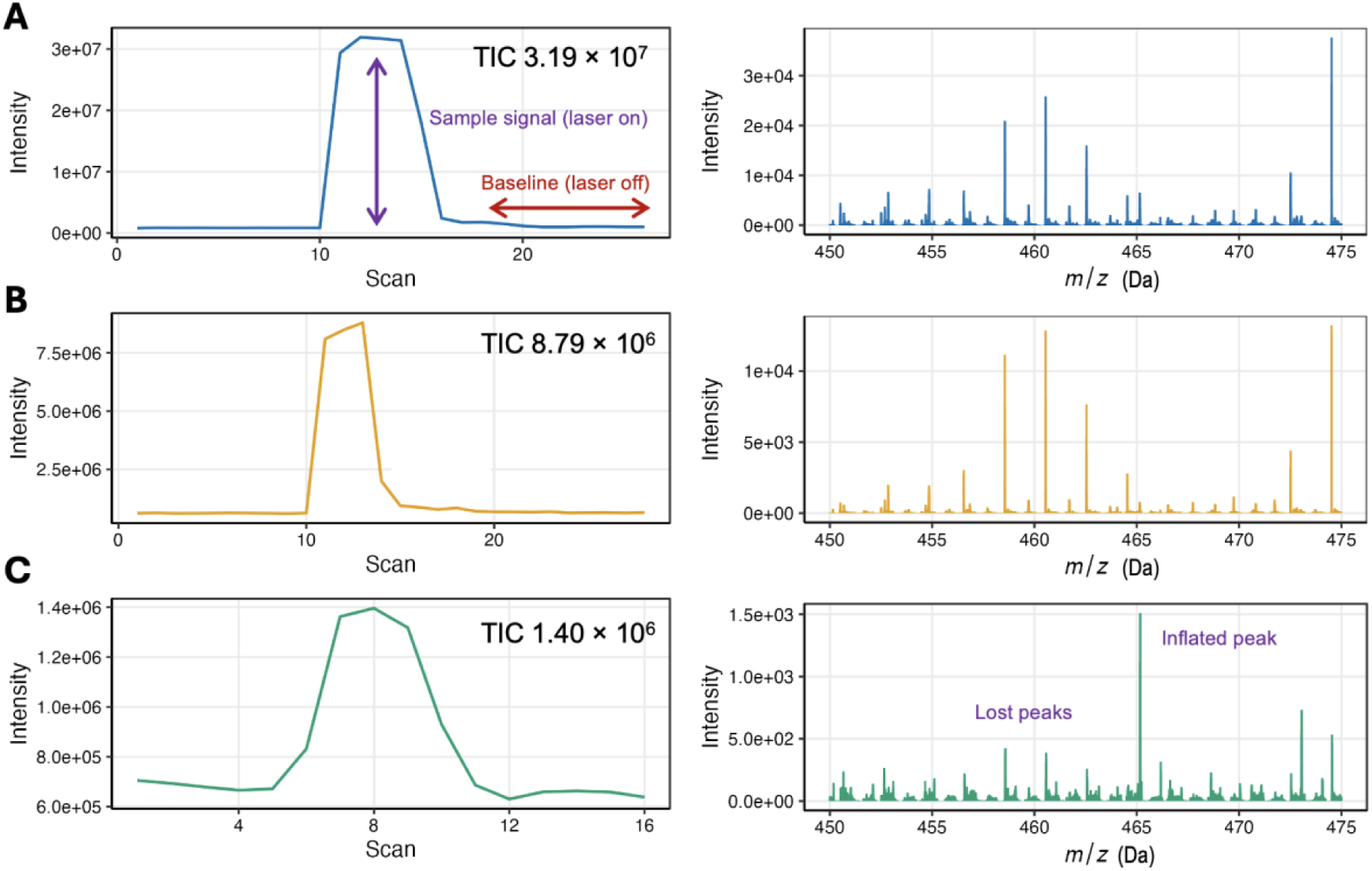
Comparison of the spectral signal of pooled urine following LA-REIMS analysis under different acquisition modes. The total ion current (TIC) of the pool is presented left, specific molecular features in the spectra are presented right: **(A)** ToF-only acquisition (ion mobility bypassed), **(B)** optimized IMS transmission conditions and **(C)** default cIMS settings.

To mitigate signal loss, optimization efforts focused on reducing the total cIMS duty cycle while maintaining sufficient ion mobility separation and avoiding wrap-around artifacts^28,29^. Shortening the total cycle time from default 51.6 to 37.8 ms and introducing a traveling-wave (TW) voltage ramp during both the mobility separation and ion ejection phases, described previously as signal temporal compression effect^30^, resulted in substantial signal recovery. Under these conditions, the average TIC increased to 1.23 × 10⁷ ± 1.90 × 10⁵ (Fig. 1B), representing a near order-of-magnitude improvement relative to default cIMS operation. Moreover, the relative abundance of individual peaks in the averaged untargeted urine fingerprint was comparable to that observed in ToF-only acquisition (Fig. 1C), indicating that the optimized cIMS settings restored signal transmission while largely preserving spectral profiles.

Following establishment of a shortened, wrap-around-free cycle, a design-of-experiments (DoE) approach was applied to evaluate the influence of TWIMS parameters on analytical performance (SI Note 2). Effect tests showed that the TW ramping rate was the primary determinant of total feature count (F = 4.59, p = 0.043), whereas TW velocity exhibited a weaker, non-significant effect (F = 3.18, p = 0.088). The lowest TW height ramping rate achievable by the instrument (1 V/ms) produced the highest number of detected untargeted features within the tested voltage bounds. This observation is consistent with TW height governing ion transmission efficiency across broad ion mobility distributions^31^. Repeatability analyses reinforced this trend, with TW ramping rate showing the strongest overall model impact (F = 35.61, p < 0.0001). Although increasing TW velocity (from 375 to 425 m/s) produced modest gains in total feature counts, these improvements did not translate into higher numbers of reproducibly detected features. Collectively, these findings establish TW ramping rate as the dominant control parameter for robust feature detection under duty-cycle restricted conditions, whereas velocity adjustments provide only marginal, non-reproducible gains (detailed parameter settings in SI Table 2).

### 2. LA-REIMS-cIMS supports selective filtering of matrix-derived background signals from raw biofluid spectra

Having established stable single-pass cIMS operating conditions, we next examined how the addition of mobility information reshapes the structure and chemical composition of untargeted LA-REIMS biofluid fingerprints. For this purpose, pooled biofluid samples (urine, saliva, and feces; 10 replicates each) were analyzed under these conditions (SI Table 2) to evaluate systematic signal organization across the *m/z* and ion mobility arrival-time dimensions. Ion mobility separation revealed distinct, recurring signal populations within the overall feature space (Fig. 2).

**Fig 2.**
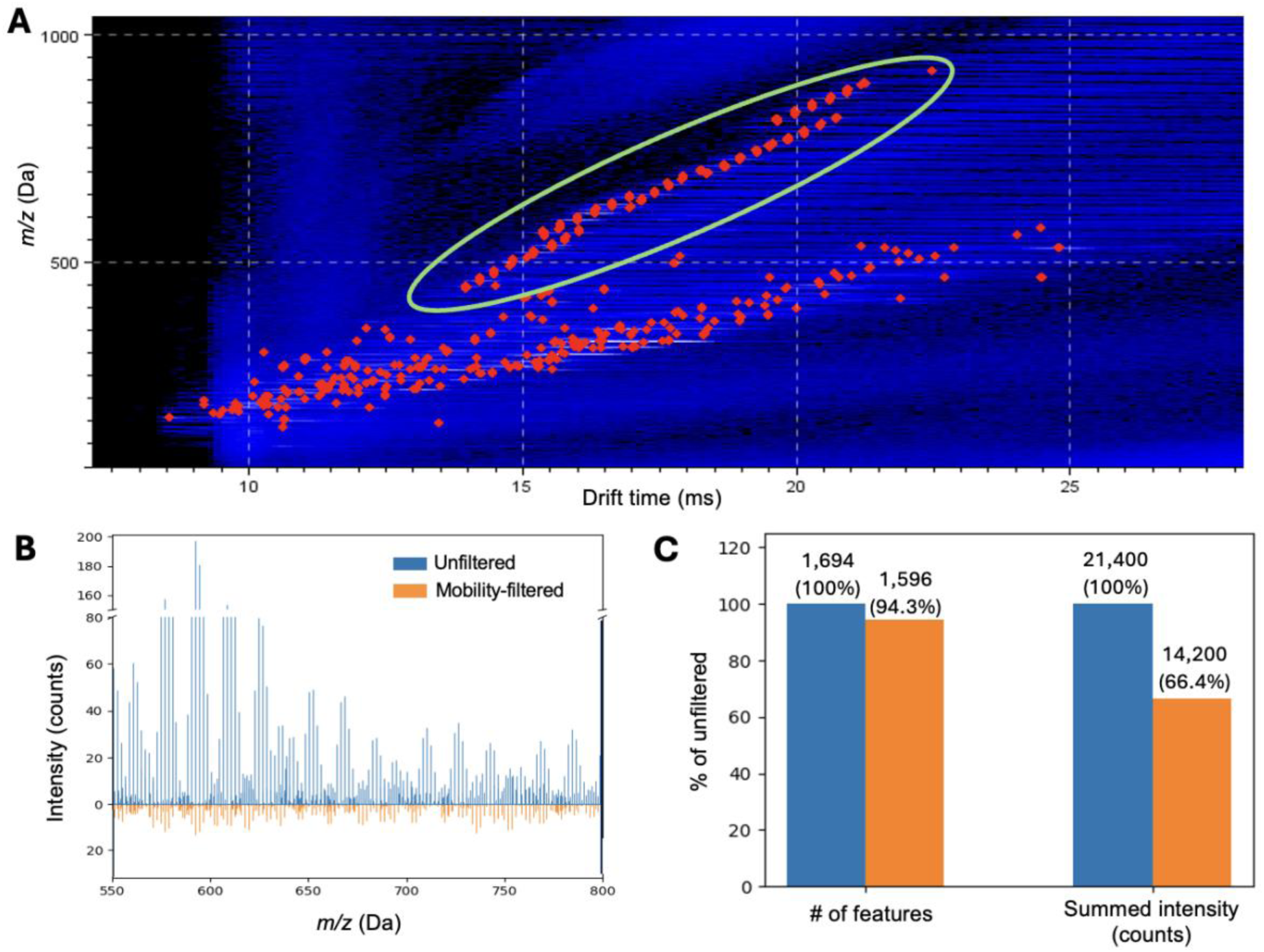
Ion mobility-resolved identification and filtering of salt-derived cluster ions in untargeted LA-REIMS-cIMS biofluid spectra. **(A)** Representative *m/z* versus ion mobility arrival-time heatmap of single pooled urine spectrum showing a primary metabolite-rich mobility band and a secondary high-density band attributed to salt-derived cluster ions (highlighted in green). **(B)** Mirrored raw-intensity spectra (550-800 *m/z*) of averaged pooled urine processed without (blue) and with mobility filtering (orange). The broken y-axis (0-80 and 150-200 intensity counts) highlights dominant salt clusters in unfiltered data and their elimination after filtering. **(C)** Feature count and summed intensity before (blue) and after mobility filtering (orange), calculated from average values and expressed relative to unfiltered spectra (100%). Feature number is largely retained (94.3%), whereas total intensity decreases (66.4%), indicating selective removal of high-intensity matrix signals.

In untargeted LA-REIMS analyses of urine and saliva, but not feces, ion mobility separation consistently revealed a secondary signal band, defined as a dense, contiguous population of features in the arrival-time space^32^, in addition to the primary metabolite-rich band (Fig. 2A). This secondary band was characterized by highly regular *m/z* spacing and arrival time patterns distinct from the endogenous metabolite signal space. Based on the observed spacing and mobility behaviour, these features were tentatively attributed to salt-derived cluster ions, which have been reported previously (in e.g. electrospray and matrix-assisted desorption ionization^33,34^) in relation to biofluids and other salt-rich matrices. The assignment of this secondary mobility band to salt-derived clusters was experimentally confirmed by analysis of a physiological sodium chloride solution under identical acquisition conditions, which reproduced the characteristic mobility pattern observed in urine and saliva samples (SI Fig. 1). Accordingly, the secondary mobility band predominantly reflects a systematic inorganic background that does not contribute to the intrinsic biochemical fingerprint of the sample and primarily reduces analytical clarity by obscuring biologically informative metabolite features.

Because the observed salt-cluster band overlaps with the mass range in which many endogenous metabolites are detected (approximately 300 to 800 Da), it can artificially inflate the number of detected spectral features and interfere with fragmentation spectra. In addition, these cluster ions are typically among the most intense species in the spectrum, thereby dominating the overall signal intensity and increasing ion competition during acquisition. Notably, filtering out this salt-cluster mobility band from urine spectra reduced the number of detected peaks by only 5.7% (from 1693.5 ± 55 to 1596.5 ± 51), whereas the summed intensity of all detected peaks decreased by 33.6% (from 2.14 × 10⁴ to 1.42 × 10⁴). Importantly, filtering out this dominant background also reduced spectral overlap and improved detectability of low-abundance organic features from the same region, without substantially altering overall feature count compared to non-filtered spectral data (Fig 2B and C). It is imperative to acknowledge though that ion mobility filtering is incapable of eradicating undesirable signals in untargeted spectra that do not demonstrate distinguishable mobility behaviour (e.g., detector-associated artifacts, SI Fig. 2). In such instances, its implementation should be accompanied by additional computational background signal removal methodologies, such as peak intensity and S/N ratio thresholds and blank subtraction^34^.

### 3. Targeted profiling of biofluid metabolome and lipidome class representatives demonstrates broad molecular coverage of LA-REIMS-cIMS

To evaluate the intrinsic molecular coverage of LA-REIMS and to assess how ion mobility integration alters compound detectability and annotation potential, a targeted panel of analytical standards representative of major metabolome and lipidome classes (n = 101, logP −5.30 to 19.40, SI Table 4) was analyzed under both ToF-only and cIMS acquisition modes.

Under ToF-only acquisition, all standards (100% of the panel) were consistently detected across replicate analyses (n = 3), demonstrating broad intrinsic molecular coverage of REIMS for chemically diverse small molecules and lipids (Fig. 3). A total of 63 compounds (62% of the panel) were observed in both ionization modes, while a subset showed polarity-specific detection (24 negative-only, 14 positive-only; SI Table 4), reflecting expected class-dependent ionization behaviour (e.g., preferential protonation of acylcarnitines^35^). Multiple adduct types were frequently observed, including chloride adducts in negative mode and alkali metal adducts positive mode, consistent with the presence of inorganic salts in untreated biological matrices^36,37^.

**Fig 3.**
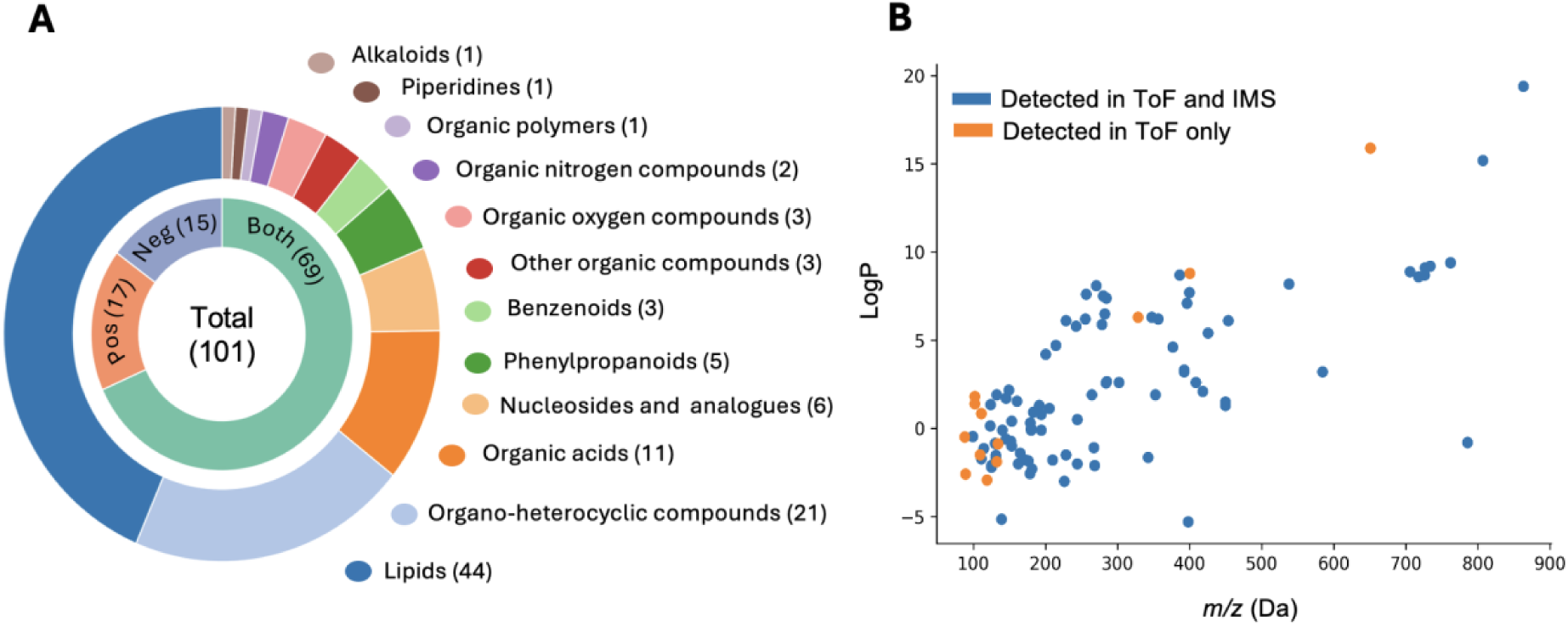
Chemical space and detection characteristics of the targeted metabolite panel following LA-REIMS-cIMS analysis. (**A**) Nested donut chart summarizing the taxonomic distribution and polarity coverage of the 101-compound targeted panel. The outer ring represents HMDB superclasses^22^, with segment size proportional to the number of compounds in each superclass (n = 101). The inner ring indicates polarity coverage. (**B**) Distribution of targeted compounds in physicochemical space plotted as *m/z* versus logP.

By comparison, UHPLC-HRMS analysis of the same analytical standard panel, performed using a Vanquish™ Duo UHPLC system coupled to an Orbitrap Exploris™ 120 HRMS (HESI-II source, polarity switching) and based on our previously validated fecal metabolomics and lipidomics methods^18^, yielded spectra dominated by protonated and deprotonated molecular ions. This difference in adduct distribution reflects effective salt removal during chromatography and differences in ionization physics rather than intrinsic differences in compound detectability^18^. Despite differences in ionization mechanisms, compound detectability showed substantial overlap between REIMS and HESI-based ionization, with no systematic compound-class-specific ionization bias observed within the tested panel and concentration range.

Introduction of cIMS under optimized conditions resulted in a reduction in compound detectability, with 85 of the 101 standards remaining detectable (Fig 3B; SI Table 5). To assess whether compound loss followed systematic trends, detectability was examined as a function of singly charged *m/z* and polarity (LogP). No clear monotonic dependence was observed across either parameter, indicating that transmission through the ion mobility device is not governed by simple univariate physicochemical cut-offs. Nonetheless, compounds at the lower extreme of the investigated *m/z* range (≤ 100 Da; e.g., valeric and isovaleric acid) and those exhibiting low signal intensities under ToF-only conditions were more frequently absent under cIMS conditions. These boundary effects suggest that transmission losses arise from multivariate ion transport and duty-cycle constraints rather than predictable linear trends. This observation is consistent with the results of previous TWIMS/cIMS studies, which indicated that ions with low *m/z* values are more prone to probabilistic transmission loss^38,39^. Importantly, however, the majority of structurally and chemically diverse standards remained detectable in cIMS, indicating that mobility integration largely preserves broad metabolome coverage at the compound class level.

Beyond detectability, the targeted standards dataset was used to evaluate the accuracy of feature descriptors underpinning untargeted analysis, namely mass accuracy and collision cross section (CCS) performance under LA-REIMS-cIMS conditions (SI Table 5). Following CCS calibration, the relative error in CCS assignment across detected standards was moderate (mean deviation 4.0%), while mass accuracy after drift-time correction remained high (mean 2.4 ppm). The observed CCS deviations are consistent with previously reported inter-laboratory and inter-platform variability and likely reflect systematic differences between literature CCS values, frequently determined using drift tube or other IMS platforms, and those measured on the cyclic TWIMS used here^13^. Although CCS accuracy alone is insufficient for low-tier compound identification in untargeted scenarios^40^, the combination of high-accuracy *m/z* measurements with reproducible CCS values provides orthogonal information that improves downstream feature matching and annotation beyond what accurate mass alone can achieve^41^. In the context of ambient LA-REIMS analysis, this combined descriptor supports class-level metabolite annotation, facilitates exclusion of false positives, and provides structural context that is otherwise inaccessible in the absence of chromatographic separation.

### 4. LA-REIMS cIMS reveals lipid organization and enhances isomer discrimination via multiple-cycle separation

Next, we examined how ion mobility separation extends the structural interpretability of untargeted LA-REIMS datasets beyond targeted compound detection and enables discrimination of closely related isomeric species within an ambient workflow. Across pooled urine samples, mobility-resolved LA-REIMS spectra revealed compact ion clusters occupying shared arrival-time domains and clearly separated from the salt-cluster band (Fig. 4A). These features, predominantly within the *m/z* range of 500-800 Da, formed coherent mobility bands consistent with lipid homologous series. Within these domains, ions sharing common backbone structures and similar total carbon numbers grouped together, while variations in acyl-chain composition produced systematic yet subtle mobility offsets^42^. Comparable lipid organization was also observed in saliva and fecal pools, indicating that mobility-resolved clustering of lipid homologues is a consistent feature across biofluid matrices.

**Fig 4.**
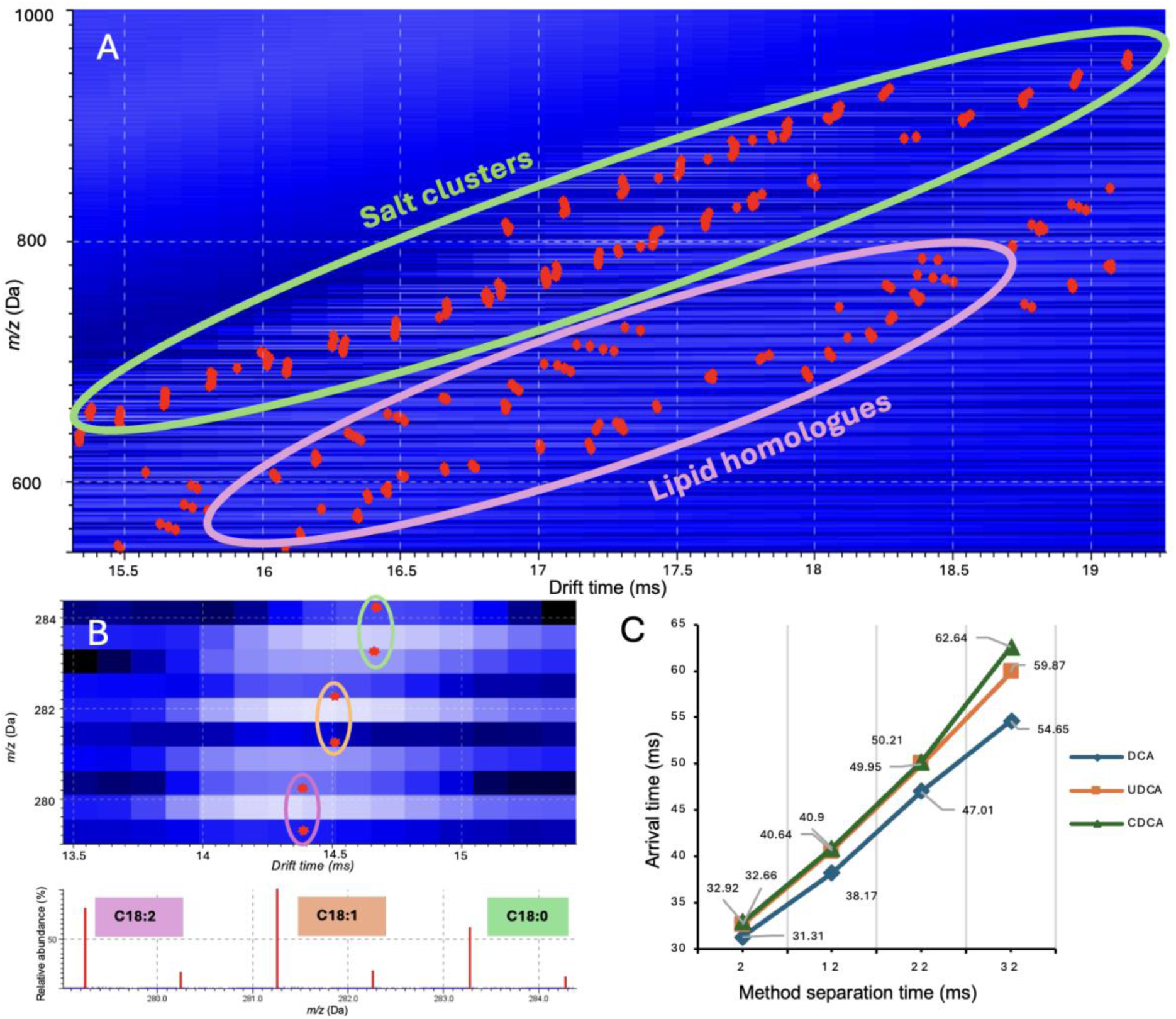
Mobility-resolved organization of lipid homologues and fatty acid saturation forms in LA-REIMS-cIMS biofluid spectra. **(A)** Pooled urine spectrum showing lipid homologues which form a distinct mobility band separated from salt-derived cluster ions. **(B)** Pooled fecal spectrum showing an analogous mobility-resolved pattern for free fatty acids, including linoleic acid (C18:2; *m/z* 279.23), oleic acid (C18:1; *m/z* 281.25), and stearic acid (C18:0; *m/z* 283.26), along with their 13C isotopic peaks and corresponding mass spectrum. (**C**) Arrival times of the bile acid isomers deoxycholic acid (DCA), ursodeoxycholic acid (UDCA), and chenodeoxycholic acid (CDCA) plotted as a function of effective cyclic IMS separation time (2-32 ms).

To illustrate this behaviour in a chemically simplified system, fecal free fatty acids were investigated more in depth (Fig. 4B). Unlike complex glycerophospholipids, free fatty acids lack bulky headgroups and extensive adduct diversity, providing a clearer view of mobility shifts associated solely with saturation state. Adjacent species differing by one double bond were separated by 2.016 Da, corresponding to the mass difference of two hydrogen losses^17,42^. Under single-pass cIMS conditions, each additional double bond induced a reproducible drift-time shift of approximately 0.13 ms in this fatty acid series. In this framework, the core molecular structure defines the absolute drift-time range, while incremental changes in unsaturation induce reproducible relative shifts in drift time within a given dataset^11,43^. Together, these relationships enable biologically meaningful grouping of related lipid features directly from intact LA-REIMS-cIMS fingerprints, providing structurally information.

Having demonstrated that ion mobility separation adds structurally meaningful organization to untargeted LA-REIMS fingerprints, we next evaluated the extent to which such separation can be leveraged for explicit isomer discrimination under practical acquisition constraints. In the subsequent targeted experiments, analytical standard solutions of secondary bile acids, their glycine- and taurine-conjugated forms, and phosphocholine (PC) lipids were examined as model compounds to assess the performance of the established LA-REIMS-cIMS workflow to generate isomer-resolved chemical signatures under both short and extended separation conditions. The selected analytes comprise structurally similar isomers, are abundant in fecal samples, and are biologically relevant^44,45^, making them suitable benchmarks for assessing cIMS performance in the context of biofluid analysis.

For bile acids, multiple ion forms were detected, including deprotonated monomers, hydrogen-bonded dimers, sodium-bound dimers, and, when their *m/z* values fell within the acquisition mass range, hydrogen-bonded trimers (Table 1), consistent with the structural complexity and adduct diversity observed in bile acid research^46^. While individual standards exhibited distinct arrival times, this separation did not translate to isomer resolution in mixtures under single-cycle IMS conditions. Arrival-time differences between deprotonated monomers were small (<5 ms) and insufficient to resolve isomeric bile acids based on their [M-H]⁻ ions, irrespective of conjugation state. In contrast, partial isomer discrimination was observed for hydrogen-bonded dimers. Under single-cycle IMS conditions, the chenodeoxycholic acid (CDCA)-dominated hydrogen-bonded dimer exhibited higher mobility than other bile acid dimers, with an arrival time difference of 5.08 ms relative to nearest co-migrating dimer, enabling its resolution in mixtures. This effect diminished upon glycine conjugation and was absent for taurine-conjugated bile acids, which collapsed into a single unresolved mobility feature, reflecting the dependence of IMS separation on the ion conformation and adduct chemistry^47^. Extending the cIMS separation time enabled improved bile acid isomer resolution. In mixtures of deoxycholic acid (DCA), ursodeoxycholic acid (UDCA), and CDCA, deprotonated CDCA was resolved from the other isomers at a 12 ms separation cycle, while all three bile acids produced distinct mobility-resolved signals at 32 ms (as their [M-H]⁻ forms; Fig 4C). Although cIMS provides sufficient tunability to resolve closely related bile acid isomers, optimal separation performance is compound- and class-dependent and requires targeted optimization^48,49^.

In contrast, fully saturated positional phosphocholine isomers with mirrored acyl-chain distributions (PC 16:0/18:0 and PC 18:0/16:0) remained unresolved under all tested conditions. No separation was observed even at extended cIMS separation cycles of 100 and 200 ms (arrival times of 89.49 and 89.53 ms for the respective chlorine adducts). The minimal 0.04 ms difference is consistent with the negligible collision cross section variation expected for exchange of two linear, fully saturated acyl chains, which introduces only marginal perturbation of the overall ion conformation^50^. Notably, these species are also challenging to resolve chromatographically in routine workflows, as reversed-phase retention is dominated by bulk hydrophobicity (total carbon number and unsaturation) and HILIC retention by headgroup polarity rather than acyl-chain position^51,52^. Consequently, robust differentiation of structurally subtle positional isomers often requires complementary structural information from MS/MS-based strategies (e.g., ozone dissociation or related reaction/fragmentation methods), which can be incorporated alongside both LC-MS and IMS-MS measurements^53,54^.

### 5. Analytical scope, applications, and limitations of LA-REIMS-cIMS in ambient biofluid metabolomics

cIMS introduces multidimensionality into the chromatography-free AIMS data and can substantially improve the specificity of ambient biofluid fingerprints by providing an orthogonal separation dimension. This is particularly valuable under conditions of high chemical noise or extensive matrix interferences, and when structural resolution, such as isomer discrimination or CCS-based annotation, is central to the analytical objective. In addition, CCS-based annotation is generally more reproducible across platforms than chromatographic retention time matching, making it more amenable to standardization^13,26,41^.

However, for high-throughput metabolic fingerprinting workflows where analytical performance is driven primarily by global spectral patterns rather than feature-level annotation, MS-only acquisition is often sufficient and can offer practical advantages. In such scenarios, the added dimensionality provided by ion mobility may not always translate into proportional gains in downstream interpretability, while increased acquisition speed and sensitivity may be achieved by maximizing duty cycle. When matrix-derived artefacts are limited, or when biological variability dominates spectral differences, omitting IMS can therefore represent a pragmatic and effective strategy. In this context, ambient ionization platforms capable of operating in either IMS-enabled or MS-only mode provide flexibility to address complementary analytical needs, with the optimal acquisition strategy determined by matrix complexity and study objectives.

### 6. Conclusion

This study establishes practical conditions under which cyclic ion mobility spectrometry can be integrated with LA-REIMS for rapid, chromatography-free biofluid metabolomics despite the transient ion generation inherent to ambient ionization workflows. By implementing duty-cycle-aware optimization, single-pass cIMS operation was achieved that preserved a substantial fraction of the total ion current while remaining compatible with the temporal constraints of LA-REIMS acquisition. While the magnitude of signal recovery and separation performance depends on instrument architecture, ion source design and sample acquisition method, the operational principles described here are broadly applicable to transient ambient ionization workflows.

Furthermore, ion mobility separation reorganizes complex LA-REIMS spectra into structured mass-mobility feature domains, enabling selective suppression of matrix-derived salt cluster ions and revealing chemically coherent signal populations, particularly within lipid classes. The combination of accurate mass and reproducible collision cross-section measurements supported class-level metabolite annotation and enabled conditional discrimination of selected isomeric species, illustrating both the analytical capabilities and intrinsic limitations of cIMS under short separation times.

Collectively, these results define the analytical operating space of LA-REIMS-cIMS and clarify the trade-offs between sensitivity, throughput, and chemical specificity associated with ion mobility integration. The framework presented here provides guidance for informed selection of IMS-enabled versus MS-only acquisition strategies in ambient metabolomics applications where chromatographic separation is impractical and enhanced chemical resolution is required.

## Acknowledgments

This work was funded in part by the European Union (ERC project MeMoSA, 2023-CoG, 101124151). Views and opinions expressed are however those of the author(s) only and do not necessarily reflect those of the European Union or the European Research Council. Neither the European Union nor the granting authority can be held responsible for them. The authors acknowledge Waters Corporation for assistance with the LA-REIMS-cIMS integration and technical support.

## Author contributions

**V.P.**: Conceptualization, methodology, investigation, data curation, formal analysis, visualization, and writing - original draft; **N.V.d.V.**: Investigation, data curation, formal analysis, visualization, interpretation, reviewing and writing; **A.V.B.**: Investigation, data curation, formal analysis, and interpretation; **J.D.D.M.**: Conceptualization, reviewing and writing. **L.V.**: Conceptualization, funding acquisition, supervision, project administration, reviewing and writing. All authors contributed to manuscript revision and approved the final version.

## Data and code availability

Processed data, representative spectra, and additional figures supporting the findings of this study are provided in the Supporting Information. Raw mass spectrometry data will be made publicly available via MetaboLights repository **MTBLS2605** upon publication. The R Shiny application code for spectra curation is available via https://github.com/UGent-LIMET/MS_spectra_curation.

