## Supplemental Material for "Ion Mobility-Enhanced LA-REIMS Improves Molecular Resolution in Ambient Biofluid Metabolomics"

**Supplemental Note 1. ToF parameter optimization on SELECT SERIES Cyclic travelling wave ion mobility time-of-flight mass spectrometer coupled to LA-REIMS ablation and ionization source.**

As differences in ion optics and detector configurations did not allow direct transfer of the LA-REIMS methods optimized for the Xevo G2-XS to the SELECT SERIES Cyclic (both from Waters Corp.) a short DoE-based optimization round was performed. The following parameters were evaluated: laser Q-switch delay time (165 - 200 µs), MS cone voltage (20 - 100 V), MS heater bias voltage (10 - 30 V), solvent (isopropanol) flow rate (0.15 - 0.3 ml/min) and MS scan time (0.1 - 1 s/scan). Parameter ranges were selected based on the results of our previous optimized and validated LA-REIMS method for the analysis of biofluids[1]. A central composite design with 3 center points was applied and processed in JMP 18 Pro (SAS institute Inc.), yielding 29 randomized experimental runs. Each experimental run was performed in triplicate. Samples from children were pooled for the analysis (n = 6, 3 overweight and 3 normal weight) and treated in the same way as described in the main text. The DoE was conducted for both negative and positive ionization mode. In positive mode, optimization experiments were restricted to fecal samples, as this matrix provides the most chemically complex signal pattern in LA-REIMS analysis and was therefore considered representative for parameter optimization. Moreover, the optimized positive-mode settings were subsequently applied for measurements of the analytical standard mixture used for targeted validation, which was analyzed under the same conditions.

Untargeted molecular fingerprints were acquired using the LA-REIMS setup[1], coupled to the SELECT SERIES Cyclic instrument[2] (Waters Corp.) operating in ToF mode (i.e., bypassing the ion mobility unit). Data preprocessing was performed with Progenesis Bridge for background subtraction and alignment, followed by Progenesis QI for peak picking and normalization (Waters Corp.). For negative ionization, samples were preprocessed in batches according to scan time settings. For positive ionization, the intensity threshold in Progenesis Bridge was adjusted per run as measured intensities varied depending on several parameters. The threshold was set at approximately one-third of the TIC, ensuring exclusion of baseline signal.

Three main analytical performance metrics were modelled: repeatability, coverage and signal magnitude. For negative ionization, the modelled responses included the number and percentage of features with CV <30%, TIC (and its standard deviation) and total detected feature count. For positive ionization, where higher measurement variability was anticipated, only the number of features with CV < 30% (and 20%) and total feature count were included. A standard least-squares model (with 2nd degree interactions) was fitted to the data for each response variable. Prediction profiles were evaluated per biofluid to determine optimal values. Unless otherwise stated, reported effects correspond to the main effects estimated from least-squares models including second-order interaction terms.

In negative ionization mode, Q-switch delay time was the most influential parameter, particularly for coverage and TIC (TIC: $\beta = -58796.26$ (linear); coverage: $\beta = -410.15$ (linear), $\beta = -375.74$ (quadratic); all $p < 0.0001$). Only in urine was heater bias voltage more predictive (coverage: $\beta = 94.81$, $p = 0.0020$; reproducibility: $\beta = 142.89$, $p = 0.0146$). In fecal samples, Q-switch delay time showed a clear optimum around 175 µs, after which TIC and feature count decreased (ΔTIC = 52%; Δ coverage = 20%). Predictions for reproducibility were generally less certain, as indicated by larger confidence intervals. Cone and heater bias voltage showed minimal effects (Δcoverage < 2%, ΔTIC < 10%, Δreproducibility < 20%). Increasing scan time improved repeatability up to approximately 0.5 s/scan, after which the effect plateaued (Δreproducibility < 10%). Flow rate had a modest effect on repeatability, with a decline beyond 0.225 ml/min, while improvement was observed up to approximately 0.2 ml/min (Δreproducibility = 11%).

Given that Q-switch delay time demonstrated the strongest and most well-defined optimum, the remaining parameters were selected to support this setting. The resulting method provided high feature counts and good reproducibility, with increased TIC variability (SD TIC of 5e4, mean TIC of 2e7), which can be mitigated through normalization. For saliva and urine, parameter values were also selected based on their effect sizes.

In positive ionization mode, effect sizes were generally smaller and uncertainty intervals wider. In fecal samples, cone and heater bias voltages, in combination with scan time, were the most influential parameters (for *cone voltage x heater bias voltage*: $\beta > 200$ and $p < 0.05$, Δcoverage < 15%, Δrepeatability = 21%; for *scan time*: Δcoverage = 19%, Δrepeatability = 42 %).

In Supplemental Table 1, an overview of optimized parameters can be found. The optimized ToF parameters established here were used as the baseline for subsequent cIMS integration and duty-cycle optimization described in Supplemental Note 2.

**Supplemental Note 2. Ion mobility parameter optimization and design-of-experiments analysis**

Cyclic ion mobility spectrometry (cIMS) parameter optimization was performed to minimize duty-cycle-related ion losses while maintaining sufficient separation performance under LA-REIMS conditions. Initial experiments employed manufacturer-recommended default cIMS parameters. Under these conditions, partial truncation of the ion arrival-time distribution and occasional wrap-around artifacts were observed, indicating incomplete capture of the transient ion population generated during laser ablation.

Optimization efforts therefore focused on reducing the total cIMS cycle time while aligning traveling-wave (TW) parameters with the arrival-time distribution of the sample ion population. The cycle time was progressively shortened while monitoring the total ion current (TIC) distribution across the arrival-time window. TW starting and ending voltages were iteratively adjusted, and TW voltage ramping was applied during both the separation and ion ejection phases to promote temporal compression of ion packets and minimize ion loss. Optimal conditions were defined by complete containment of the TIC within a single mobility cycle, with no evidence of truncation at the cycle boundaries or wrap-around into subsequent cycles. Although absolute voltage values were instrument-specific, this optimization strategy proved robust across replicate measurements and independent sample aliquots. Final optimized cIMS and TWIMS parameters used throughout the study are summarized in Supplemental Table S2.

Following establishment of a shortened, wrap-around-free cIMS cycle, a design-of-experiments (DoE) approach was applied to quantitatively evaluate the influence of key TWIMS parameters on analytical performance. A central composite design incorporating three center points was used to investigate the effects of TW velocity (375 - 425 m/s) and TW voltage ramping rate (1 - 4 V/ms). Each experimental condition was analyzed in triplicate using independent aliquots of pooled urine samples. Analytical performance was assessed using two response variables: total detected feature count and the number of reproducibly detected features, defined as features exhibiting a coefficient of variation (CV) below 30% across replicate measurements. Feature detection was performed following TIC normalization (Supplemental Fig. 3).

For total feature count, effect tests identified TW voltage ramping rate as the only statistically significant factor ($F = 4.59$, $p = 0.043$), whereas TW velocity showed a weaker and statistically non-significant effect ($F = 3.18$, $p = 0.088$). Linear regression analysis confirmed a negative effect of ramping rate on feature count ($\beta = -252$), indicating reduced feature recovery at higher ramping rates. Reproducibility was more strongly governed by TW voltage ramping rate, which exhibited a highly significant effect on the number of reproducibly detected features ($F = 35.61$, $p < 0.0001$). In contrast, TW velocity had no measurable influence on reproducibility ($F = 0.24$, $p = 0.63$). These results indicate that TW ramping rate primarily controls measurement stability under duty-cycle-limited conditions, whereas improvements associated with increased TW velocity were not consistent across replicate analyses.

To assess the robustness of these trends with respect to data processing workflows, untargeted feature detection was performed using both MS-DIAL and Progenesis QI. Absolute feature counts differed between platforms due to differences in peak-picking algorithms, noise filtering, and deconvolution strategies. However, the directionality of parameter effects, particularly the dominant influence of TW ramping rate on reproducibility, was consistent across both software tools, supporting the conclusion that the observed optimization trends reflect intrinsic analytical behavior rather than software-specific artefacts. This behavior is consistent with TW ramping rate primarily governing temporal ion packet compression under short duty-cycle conditions. In contrast, velocity-driven gains depend more strongly on ion population stability and therefore show reduced robustness across replicate measurements.

**Supplemental Note 3. Interactive R Shiny application for targeted MS peak curation.**

A custom R Shiny application was developed to support manual curation of peaks detected in standard mixtures. The application provides standardized visualization of MS spectra for candidate peaks together with background signals, allowing rapid evaluation of peak quality across positive and negative ionization modes. Prior to launching the application, raw MS data are converted to a simplified spectral format using a preprocessing script. This step imports .mzML files, performs scan selection based on the total ion chromatogram, optionally applies a mass-shift correction, and generates summed spectra for both sample and background data. These spectra are then visualized in the Shiny interface together with expected *m/z* values from an adduct library. Within the application, users evaluate candidate peaks by comparing signal intensity, background signal, mass accuracy, and signal-to-noise ratio. The curated results are exported as a table containing compound and adduct identifiers together with peak metrics and the assigned classification.

The complete source code and example data are available at GitHub: https://github.com/UGent-LIMET/MS_spectra_curation.

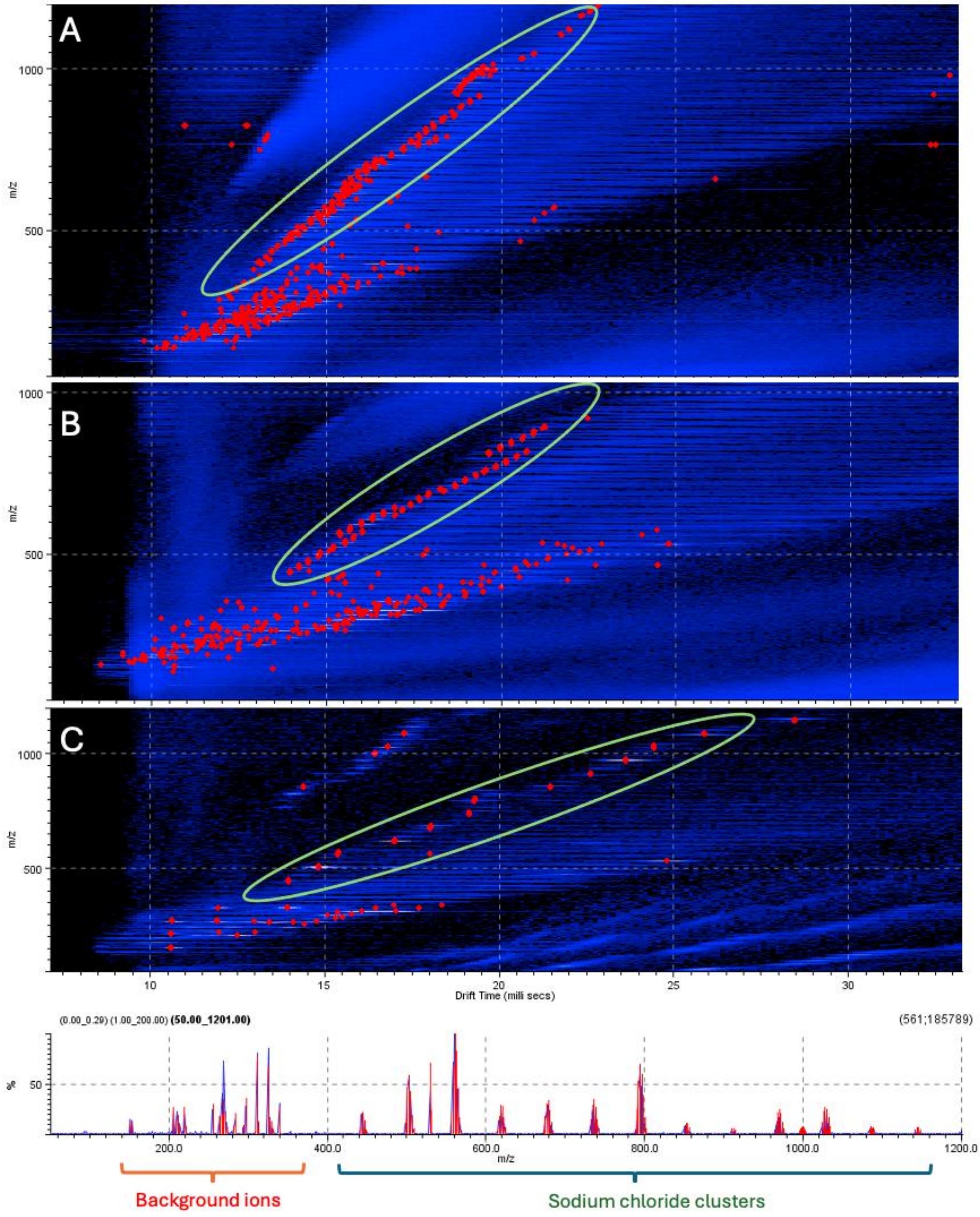


**Supplemental Fig 1. LA-REIMS-cIMS spectra of saliva (A), urine (B), and 0.9% sodium chloride solution (C, with corresponding mass spectrum).** In all spectra, a repetitive pattern in the upper mobility band (highlighted by the green oval) corresponds to signals arising from salt cluster ions. These clusters occupy a defined arrival-time region within the combined m/z-drift time space. During downstream data processing, signals within this region were selectively excluded, enabling clearer identification of the underlying biofluid molecular fingerprint.

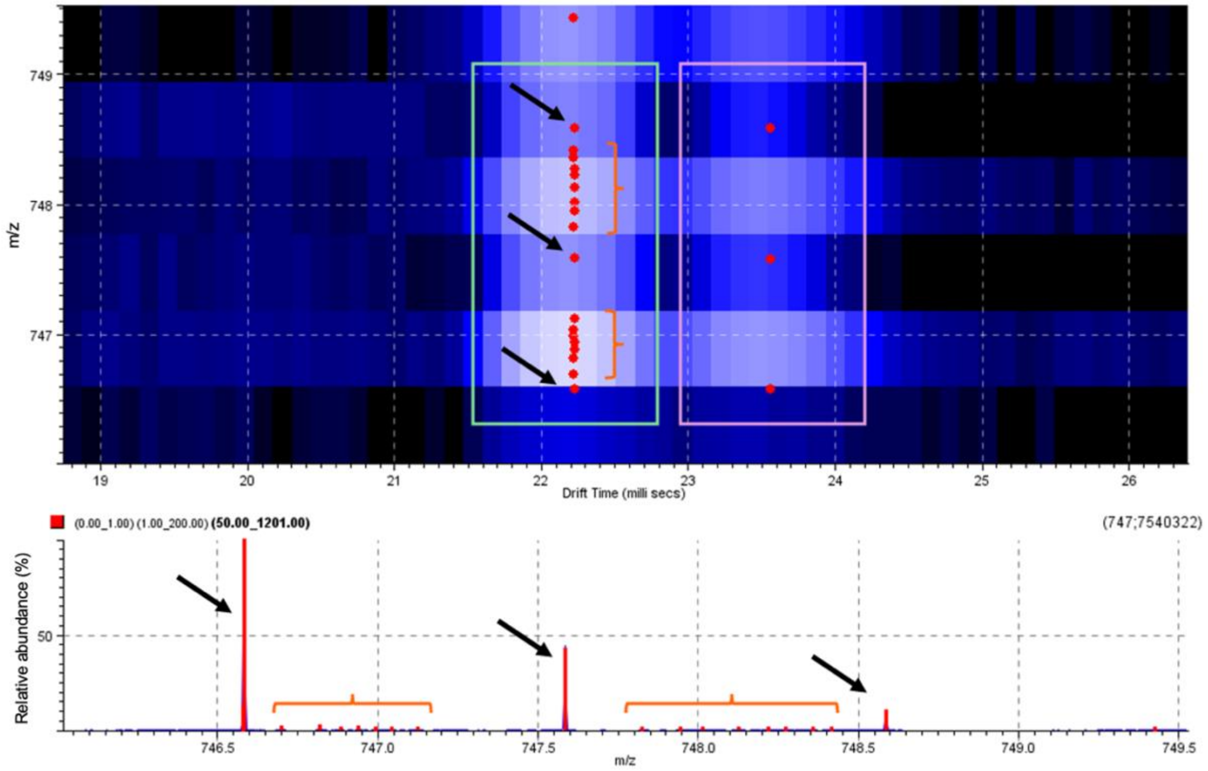


**Supplemental Fig 2. Two-dimensional *m/z*-arrival time plot and corresponding mass spectrum of the analytical standard 1-palmitoyl-2-stearoyl-*sn*-glycero-3-phosphocholine (PC 16:0/18:0), detected as the demethylated ion at *m/z* 746.55.** The dominant peaks (indicated by black arrows and green box) correspond to the isotopic distribution of the compound. Lower-intensity peaks following the main isotopic signals arise from detector oscillation artifacts, with regions containing multiple artifact peaks highlighted by bracketed regions. As these artifact peaks originate from the true compound signal and are generated at the detector level, they exhibit identical arrival times (ion mobility behavior) to the parent ions and therefore fall within the organic molecule mobility band, rendering them unaffected by band-based signal filtering. Notably, a subset of peaks corresponding to demethylated compound arises from post-mobility fragmentation of the $[M + Cl]^-$ adduct of PC 16:0/18:0 at *m/z* 796.55, appearing in the two-dimensional plot at later arrival times (signals in pink box on the plot). These peaks are of substantially lower intensity, and no associated artifact signals are observed, indicating that the formation of detector oscillation artifacts is strongly dependent on signal intensity.

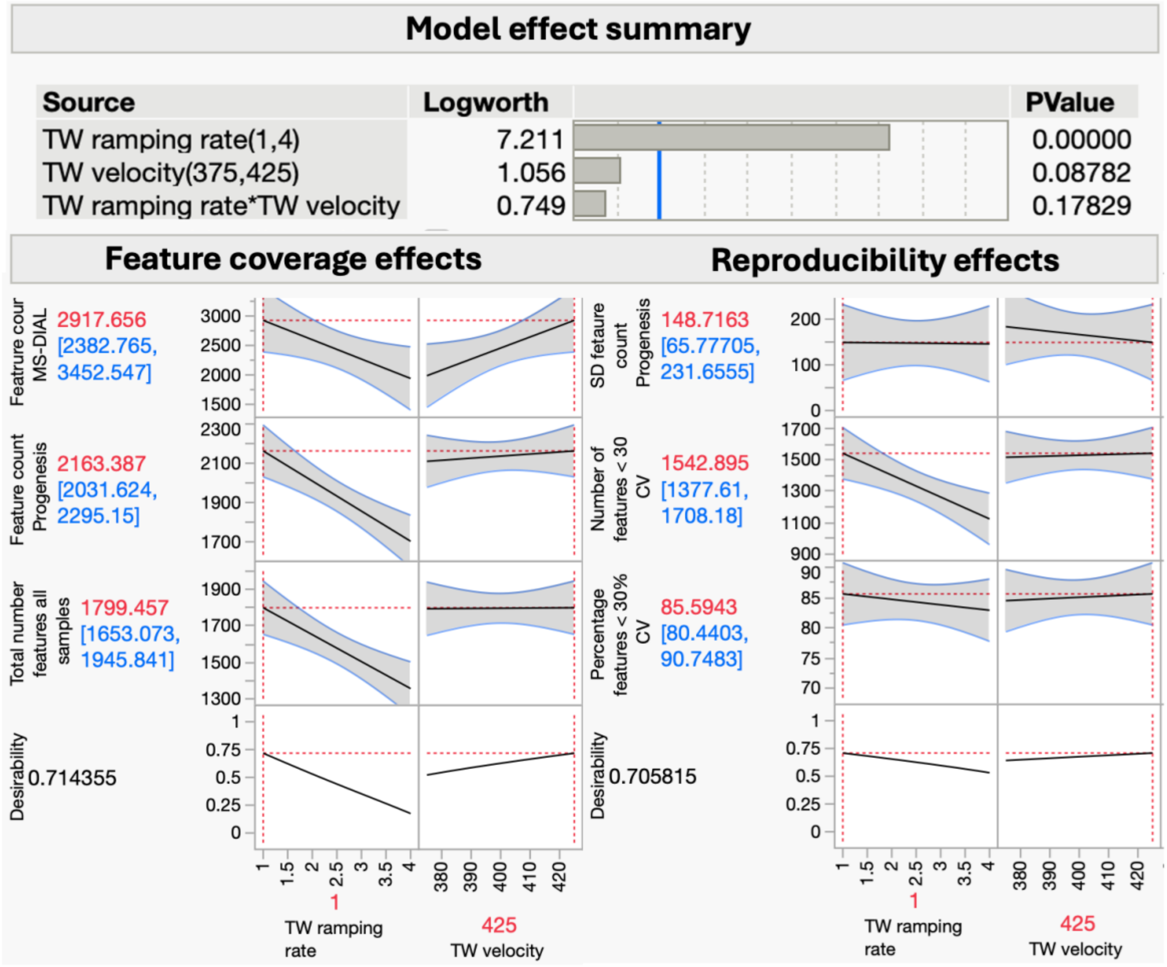


**Supplemental Fig 3. Design-of-experiments (DOE) model effects of traveling wave (TW) ramping rate and TW velocity on ion mobility performance in LA-REIMS analysis of pooled urine samples.** Effect directions for both the total number of detected features and the number of reproducibly detected features (CV < 30%, n = 3) were primarily governed by the TW ramping rate rather than TW velocity. Lower ramping rates resulted in increased feature detection and, consequently, a higher number of reproducibly detected features, independent of the data processing platform (Progenesis QI and MS-DIAL). Higher TW velocities showed a modest positive trend; however, no statistically significant effect was observed.

**Supplemental Table 1. Overview of optimized source, sampling, and ToF acquisition parameters used for measurements on the SELECT SERIES™ Cyclic™ IMS instrument.**

| Parameter | Negative | | | Positive |
|---|---|---|---|---|
| | Saliva | Feces | Urine | Feces |
| Q-switch (µs) | 180 | 175 | 180 | 180 |
| Cone voltage (V) | 100 | 80 | 100 | 60 |
| Heater bias voltage (V) | 30 | 30 | 30 | 20 |
| Flow rate (mL/min) | 0.2 | 0.2 | 0.2 | 0.2 |
| Scan time (s/scan) | 0.5 | 0.5 | 0.5 | 0.5 |

**Supplemental Table 2. Overview of optimized parameters for operating the SELECT SERIES™ Cyclic™ IMS instrument in mobility mode.**

| Parameter | Value |
|---|---|
| Pushes per bin | 2 |
| TW velocity (m/s) | 375 |
| TW start height (V) | 15 |
| TW limit height (V) | 20 |
| TW ramping rate (V/ms) | 1 |
| ADC start delay (ms) | 10 |
| IMS cycle time (ms) | 33.4 |

**Supplemental Table 3. Targeted compounds used for CCS calibration.**

| Compound | Formula | *m/z* | Adduct | Ref. CCS (Å²) |
|---|---|---|---|---|
| **Negative polarity** | | | | |
| 2-Aminoethylphosphonate | C2H8NO3P | 124.0164 | [M-H]- | 120.3 |
| L-Isoleucine | C6H13NO2 | 130.0868 | [M-H]- | 130.6 |
| Dopamine | C8H11NO2 | 152.0712 | [M-H]- | 133.3 |
| L-Phenylalanine | C9H11NO2 | 164.0712 | [M-H]- | 140.1 |
| 1-Methyl-6,7-dihydroxy-1,2,3,4-tetrahydroisoquinoline | C10H13NO2 | 178.0868 | [M-H]- | 139.7 |
| Abscisic Acid | C15H20O4 | 263.1284 | [M-H]- | 173.8 |
| Inosine | C10H12N4O5 | 267.0730 | [M-H]- | 159.6 |
| Deoxycholic acid | C24H40O4 | 391.2849 | [M-H]- | 202.1 |
| **Positive polarity** | | | | |
| L-Threonine | C4H9NO3 | 120.0660 | [M+H]+ | 127.3 |
| 3-Hydroxyanthranilic acid | C7H7NO3 | 154.0504 | [M+H]+ | 131.3 |
| L-Tyrosine | C9H11NO3 | 182.0817 | [M+H]+ | 145.5 |
| L-Tyrosine | C9H11NO3 | 204.0630 | [M+Na]+ | 143.2 |
| 5-Hydroxyindole-3-acetic acid | C10H9NO3 | 192.066 | [M+H]+ | 141.1 |
| Xanthurenic acid | C10H7NO4 | 228.0273 | [M+Na]+ | 148.0 |
| Inosine | C10H12N4O5 | 267.073 | [M+Na]+ | 159.6 |
| Piperine | C17H19NO3 | 286.1438 | [M+H]+ | 171.4 |
| Piperine | C17H19NO3 | 308.1263 | [M+Na]+ | 166.9 |
| 1-Palmitoyl-glycero-3-phosphoethanolamine (16:0) | C21H44NO7P | 454.2928 | [M+H]+ | 213.9 |

**Supplemental Table 4. Targeted panel composition with minimal detected concentration, adduct forms and polarity in the ToF-mode of LA-REIMS-cIMS setup.**

| Compound | Superclass (HMDB) | Class (HMDB) | LogP | Min. detected concentration (ng/µL) | Detected adducts |
|---|---|---|---|---|---|
| 1-Methyl-6,7-dihydroxy-1,2,3,4-tetrahydroisoquinoline | Organoheterocyclic compounds | Tetrahydroisoquinolines | 0.3 | 10 | [M+H]+; [M-H]- |
| 1-Methyl-hydantoin | Organoheterocyclic compounds | Azolines | -1.15 | 10 | [M-H]- |
| 1-Oleoyl-rac-glycerol (MG(18:1(9Z)/0:0/0:0)) | Lipids and lipid-like molecules | Glycerolipids | 6.2 | 10 | [M+Cl]-; [M+Na]+ |
| 1-palmitoyl-2-myristoyl-sn-glycero-3-phosphocholine (= PC(16:0/14:0) | Lipids and lipid-like molecules | Glycerophospholipids | 8.9 | 10 | [M+Cl]-; [M+H]+; [M+Na]+; [M-H]- |
| 1-Palmitoyl-2-oleoyl-glycero-3-phosphoethanolamine (16:0/18:1) (= PE( 16:0/18:1) | Lipids and lipid-like molecules | Glycerophospholipids | 8.6 | 10 | [M+H]+; [M+Na]+; [M-H]- |
| 1-Palmitoyl-2-stearoyl-glycero-3-phosphocholine (16:0/18:0) | Lipids and lipid-like molecules | Glycerophospholipids | 9.4 | 10 | [M+Cl]-; [M+H]+; [M+Na]+ |
| 1-Palmitoyl-glycero-3-phosphoethanolamine (16:0) | Lipids and lipid-like molecules | Glycerophospholipids | 6.1 | 10 | [M+H]+; [M+Na]+; [M-H]- |
| 1-Stearoyl-2-palmitoyl-glycero-3-phosphocholine (18:0/16:0) (= PC(18:0/16:0)) | Lipids and lipid-like molecules | Glycerophospholipids | 9.4 | 10 | [M+Cl]-; [M+H]+; [M+Na]+ |
| 1,2-Dipalmitoyl-glycero-3-phosphocholine (16:0/16:0) (DPPC) | Lipids and lipid-like molecules | Glycerophospholipids | 9.2 | 10 | [M+Cl]-; [M+H]+; [M+Na]+; [M-H]- |
| 1,2-Distearoyl-sn-glycero-3-phosphate (sodium salt) | Lipids and lipid-like molecules | Glycerophospholipids | 9.1 | 10 | [M+Cl]-; [M+H]+; [M+Na]+ |
| 1,6-Anhydro-b-glucose | Organoheterocyclic compounds | Oxepanes | -2 | 10 | [M-H]- |
| 2-Aminoethylphosphonate | Organic oxygen compounds | Organooxygen compounds | -2.1 | 100 | [2M+H]+; [2M+Na]+; [M+Cl]-; [M+H]+; [M-H]- |
| 2-Deoxyuridine | Nucleosides, nucleotides, and analogues | Pyrimidine nucleotides | -1.51 | 10 | [M+Cl]-; [M+Na]+; [M-H]- |
| 2-Piperidione | Piperidines | Piperidinones | -0.46 | 10 | [M+H]+; [M+Na]+ |
| 3-Hydroxy-4-methoxycinnamic acid | Phenylpropanoids and polyketides | Cinnamic acids and derivatives | 0.79 | 10 | [M-H]- |

| | | | | | |
|---|---|---|---|---|---|
| 3-Hydroxyanthranilic acid | Benzenoids | Benzene and substituted derivatives | 0.4 | 10 | [M+H]+; [M-H]- |
| 3-Phenylpropionate | Phenylpropanoids and polyketides | Phenylpropanoic acids | 2.17 | 10 | [M+H]+; [M+Na]+; [M-H]- |
| 3-Ureidopropionate | Organic acids and derivatives | Organic carbonic acids and derivatives | -1.9 | 100 | [M+2Na-H]+; [M+Na]+; [M-H]- |
| 4-Methylcatechol | Benzenoids | Phenols | 1.34 | 10 | [M+Hac-H]-; [M+NH4]+; [M-H]- |
| 5-Hydroxyindole-3-acetic acid | Organoheterocyclic compounds | Indole-3-acetic acid derivatives | 1.3 | 10 | [M-H]- |
| Abscisic acid | Lipids and lipid-like molecules | Prenol lipids | 1.9 | 10 | [M+H]+; [M-H]- |
| Adenosine (Adenosyl) | Nucleosides, nucleotides, and analogues | Purine nucleotides | -1.1 | 10 | [M+Cl]-; [M+H]+; [M-H]- |
| alfa-Linolenic acid | Lipids and lipid-like molecules | Fatty Acyls | 5.9 | 10 | [M+H]+; [M-H]- |
| Anandamide C20:4 (AEA) | Organic nitrogen compounds | Organonitrogen compounds | 6.3 | 10 | [M+Cl]-; [M+H]+; [M+Na]+; [M-H]- |
| Bilirubin | Organoheterocyclic compounds | Tetrapyrroles and derivatives | 3.22 | 10 | [M+H]+; [M+Na]+; [M-H]- |
| Caffeine | Organoheterocyclic compounds | Imidazopyrimidines | -0.1 | 10 | [M+Cl]-; [M+H]+; [M-H]- |
| Campesterol | Lipids and lipid-like molecules | Steroids and steroid derivatives | 8.8 | 10 | [M+H]+ |
| Chenodeoxycholic acid (CDCA) | Lipids and lipid-like molecules | Bile acids, alcohols and derivatives | 3.2 | 10 | [M+Cl]-; [M+Na]+; [M-H]- |
| Cholesterol | Lipids and lipid-like molecules | Steroids and steroid derivatives | 8.7 | 10 | [M+H]+ |
| Cholesteryl linoleate | Lipids and lipid-like molecules | Steroids and steroid derivatives | 15.9 | 100 | [M+Na]+ |
| Cholic acid (CA) | Lipids and lipid-like molecules | Bile acids, alcohols and derivatives | 2.6 | 10 | [M+Cl]-; [M+H]+; [M+Na]+; [M-H]- |
| Choline chloride | Organic acids and derivatives | Organic sulfuric acids and derivatives | -5.16 | 10 | [M+Cl]- |
| Cinnamaldehyde | Phenylpropanoids and polyketides | Cinnamaldehydes | 1.9 | 10 | [2M+Na]+; [M+H]+; [M+Na]+ |
| Coenzyme Q10 | Lipids and lipid-like molecules | Prenol lipids | 19.4 | 10 | [M+H]+; [M+Na]+ |

| | | | | | |
|---|---|---|---|---|---|
| Cytosine | Organoheterocyclic compounds | Diazines | -1.73 | 10 | [M+H]+; [M+Na]+ |
| D-biotin | Organoheterocyclic compounds | Biotin and derivatives | 0.5 | 10 | [M+2Na-H]+; [M+Na-2H]-; [M+Na]+; [M-H]- |
| Deoxycholic acid (DCA) | Lipids and lipid-like molecules | Bile acids, alcohols and derivatives | 3.3 | 10 | [M+Cl]-; [M+Na]+; [M-H]- |
| Dodecanoic acid (Lauric acid) | Lipids and lipid-like molecules | Fatty Acyls | 4.2 | 10 | [M-H]- |
| Dopamine hydrochloride | Benzenoids | Phenols | -1 | 10 | [M+Cl]-; [M+Na]+; [M-H]- |
| Ergocalciferol (vitamine D2) | Lipids and lipid-like molecules | Steroids and steroid derivatives | 7.1 | 10 | [M+H]+ |
| Flavin adenine dinucleotide | Nucleosides, nucleotides, and analogues | Flavin nucleotides | -0.8 | 10 | [M+Na]+; [M-H]- |
| Galactaric acid | Organic oxygen compounds | Organic oxygen compounds | -1.8 | 10 | [M+Na]+; [M-H]- |
| Ganglioside GM3 (d18:1/16:0) | Lipids and lipid-like molecules | Sphingolipids | 12.3* | 10 | [M-H]- |
| Ganglioside GM3 (d18:1/18:0) | Lipids and lipid-like molecules | Sphingolipids | 12.3* | 10 | [M-H]- |
| Glyceryl tripalmitate (Tripalmitin) | Lipids and lipid-like molecules | Glycerolipids | 15.2 | 100 | [M+K]+; [M+Na]+ |
| Glycitein | Phenylpropanoids and polyketides | Isoflavonoids | 2.57 | 10 | [M+H]+; [M+Na]+; [M-H]- |
| Glycochenodeoxycholate (GCDCA) | Lipids and lipid-like molecules | Bile acids, alcohols and derivatives | 1.3 | 10 | [M+Cl]-; [M+H]+; [M+Na]+ |
| Glycodeoxycholate (GDCA) | Lipids and lipid-like molecules | Bile acids, alcohols and derivatives | 1.5 | 10 | [M+Cl]-; [M+H]+; [M+Na]+ |
| Hesperetin | Phenylpropanoids and polyketides | Flavonoids | 2.6 | 10 | [M+Cl]-; [M-H]- |
| Hexadecanamide (Palmitamide) | Lipids and lipid-like molecules | Fatty Acyls | 6.2 | 10 | [M+H]+; [M+Na]+ |
| Hypotaurine | Organic acids and derivatives | Sulfinic acids and derivatives | -1.5 | 100 | [M+2Na-H]+; [M+H]+ |
| Imidazolepropionic acid | Organoheterocyclic compounds | Azoles | -0.09 | 10 | [M+Cl]-; [M-H]- |
| Indole-3-carboxaldehyde | Organoheterocyclic compounds | Indoles and derivatives | 1.7 | 10 | [M+H]+; [M-H]- |

| | | | | | |
|---|---|---|---|---|---|
| Inosine | Nucleosides, nucleotides, and analogues | Purine nucleosides | -2.1 | 10 | [M+Na]+; [M-H]- |
| Isovaleric acid (= 3-methylbutanoic acid) | Lipids and lipid-like molecules | Fatty Acyls | 1.8 | 10 | [M+2Na-H]+; [M-H]- |
| L-Alanine | Organic compounds | Organic acids and derivatives | -2.6 | 10 | [M-H]- |
| L-ascorbic acid | Organoheterocyclic compounds | Dihydrofurans | -1.85 | 100 | [M+2Na-H]+; [M+Na-2H]-; [M-H]- |
| L-Carnosine | Organic acids and derivatives | Peptidomimetics | -3 | 10 | [M+Cl]-; [M+Na]+; [M-H]- |
| L-gulonic gamma-lactone | Organoheterocyclic compounds | Lactones | -2.57 | 10 | [M+Cl]-; [M-H]- |
| L-Isoleucine | Organic acids and derivatives | Carboxylic acids and derivatives | -1.7 | 10 | [M-H]- |
| L-Leucine | Organic acids and derivatives | Carboxylic acids and derivatives | -1.52 | 10 | [M-H]- |
| L-Phenylalanine | Organic compounds | Organic acids and derivatives | -1.4 | 10 | [M+Cl]-; [M+H]+; [M-H]- |
| L-Threonine | Organic acids and derivatives | Carboxylic acids and derivatives | -2.94 | 10 | [M+Cl]- |
| L-Tyrosine | Organic compounds | Organic acids and derivatives | -2.3 | 10 | [M+Na]+; [M-H]- |
| Linoleic acid | Lipids and lipid-like molecules | Fatty Acyls | 7.5 | 10 | [M+H]+; [M+Na]+ |
| Lithocholic acid (LCA) | Lipids and lipid-like molecules | Bile acids, alcohols and derivatives | 4.6 | 10 | [M+Cl]-; [M-H]- |
| Malic acid | Organic acids and derivatives | Hydroxy acids and derivatives | -0.87 | 10 | [M+Na]+; [M-H]- |
| Margaric acid (=Heptadecanoic acid) | Lipids and lipid-like molecules | Fatty Acyls | 8.1 | 100 | [M+2Na-H]+; [M-H]- |
| Myristic acid (= 1-tetradecanoic acid) | Lipids and lipid-like molecules | Fatty Acyls | 6.1 | 100 | [M-H]- |
| N-Acetylagmatine | Organic polymers | Polypeptides | -1.8 | 10 | [M+Cl]-; [M+H]+; [M+Na]+ |
| N-Acetylputrescine | Organic acids and derivatives | Carboximidic acids and derivatives | -0.84 | 10 | [M+H]+; [M-H]- |

| | | | | | |
|---|---|---|---|---|---|
| N-oleoyl-D-erythro-sphingosine (ceramide d18:1/18:1) | Lipids and lipid-like molecules | Sphingolipids | 8.7 | 10 | [M+Cl]-; [M+H]+; [M+Na]+; [M-H]- |
| N-palmitoyl-D-erythro-sphingosine (ceramide d18:1/16:0) | Lipids and lipid-like molecules | Sphingolipids | 8.2 | 10 | [M+Cl]-; [M+H]+; [M+Na]+; [M-H]- |
| Nicotinic acid | Organoheterocyclic compounds | Pyridines and derivatives | 0.15 | 10 | [M+H]+ |
| Oleic acid | Lipids and lipid-like molecules | Fatty Acyls | 6.5 | 10 | [M+Cl]-; [M+Na]+ |
| Oleoyl-L-carnitine | Lipids and lipid-like molecules | Fatty Acyls | 5.4 | 10 | [M+Cl]-; [M+H]+; [M+Na]+ |
| Palmitic acid (= hexadecanoic acid) | Lipids and lipid-like molecules | Fatty Acyls | 7.6 | 10 | [2M-H]-; [M+Na]+; [M-H]- |
| Palmitoyl-L-carnitine | Lipids and lipid-like molecules | Fatty Acyls | 7.7 | 10 | [M+Cl]-; [M+H]+; [M+Na]+ |
| Pentadecanoic acid | Lipids and lipid-like molecules | Fatty Acyls | 5.8 | 100 | [M+2Na-H]+; [M-H]- |
| Piperine | Alkaloids and derivates | NR | 2.65 | 10 | [M+H]+; [M+Na]+ |
| Piperonyl acetone | Organoheterocyclic compounds | Benzodioxoles | 1.15 | 10 | [M-H]- |
| Pregnenolone sulfate sodium | Lipids and lipid-like molecules | Steroids and steroid derivatives | 2.1 | 10 | [M+Cl]-; [M+Na]+ |
| Prostaglandine E2 | Lipids and lipid-like molecules | Fatty Acyls | 1.9 | 10 | [M+Na]+ |
| Pseudouridine | Nucleosides, nucleotides, and analogues | Nucleoside and nucleotide analogues | -2 | 10 | [M+Cl]-; [M-H]- |
| Pyrrole-2-carboxylic acid | Organoheterocyclic compounds | Pyrroles | 0.85 | 10 | [M+2Na-H]+; [M+Na-2H]-; [M-H]- |
| Pyruvic acid | Organic acids and derivatives | Keto acids and derivatives | -0.5 | 10 | [2M-H]-; [M+2Na-H]+; [M-H]- |
| Retinol acetate | Lipids and lipid-like molecules | Prenol lipids | 6.3 | 10 | [M+NH4]+; [M+Na]+ |
| S-(5'-Adenosyl)-L-methionine | Nucleosides, nucleotides, and analogues | 5'-deoxyribonucleosides | -5.3 | 10 | [M+H]+ |
| Saccharin | Organoheterocyclic compounds | Benzothiazoles | 0.91 | 10 | [M-H]- |
| Spermidine | Organic nitrogen compounds | Organonitrogen compounds | -0.62 | 10 | [M+H]+ |

| | | | | | |
|---|---|---|---|---|---|
| Stearic acid | Lipids and lipid-like molecules | Fatty Acyls | 7.4 | 10 | [2M-H]-; [M+Cl]-; [M-H]- |
| Taurine | Organic acids and derivatives | Organic sulfonic acids and derivatives | -2.2 | 10 | [M+Cl]-; [M-H]- |
| Theobromine | Organoheterocyclic compounds | Imidazopyrimidines | -0.1 | 10 | [M+Cl]-; [M+H]+; [M-H]- |
| Theophylline | Organoheterocyclic compounds | Imidazopyrimidines | 0 | 10 | [M+Cl]-; [M-H]- |
| Trehalose | Organic oxygen compounds | Organooxygen compounds | -1.65 | 10 | [M+Cl]-; [M+Na]+; [M-H]- |
| Tricosanoyl sphingomyelin d18:1/23:0 | Lipids and lipid-like molecules | Sphingolipids | 15.1* | 10 | [M+Cl]-; [M+Na]+; [M-H]- |
| Tridecanoic acid | Lipids and lipid-like molecules | Fatty Acyls | 4.7 | 10 | [M-H]- |
| Tryptamine | Organoheterocyclic compounds | Indoles and derivatives | 1.55 | 10 | [M-H]- |
| Valeric acid | Lipids and lipid-like molecules | Fatty Acyls | 1.4 | 100 | [M+2Na-H]+; [M-H]- |
| Xanthine | Organoheterocyclic compounds | Imidazopyrimidines | -0.73 | 10 | [M+Cl]-; [M-H]- |
| Xanthurenic acid | Organoheterocyclic compounds | Quinolines and derivatives | 1.14 | 10 | [M+H]+; [M-H]- |

*predicted ChemSpider MolLogP

**Supplemental Table 5. Targeted panel composition with minimal detected concentration, adduct forms and polarity in the IMS-mode of LA-REIMS-cIMS setup.**

| Compound | Formula | Mol. weight (mono-isotopic) | Adduct | *m/z* (ref) | CCS (ref) | *m/z* (measured) | \|d*m/z*\| (ppm) | CCS (measured) | \|dCCS\| (%, average) | Ref. |
|---|---|---|---|---|---|---|---|---|---|---|
| 1-Methyl-6,7-dihydroxy-1,2,3,4-tetrahydroisoquinoline | C10H13NO2 | 179.0946 | [M-H]- | 178.0868 | 139.7 | 178.0868 | 0.00 | 138.04 | 1.19 | [3] |
| 1-Methyl-6,7-dihydroxy-1,2,3,4-tetrahydroisoquinoline | C10H13NO2 | 179.0946 | [M+H]+ | 180.1024 | 142.9 | 180.1008 | 8.88 | 137.4 | 3.85 | [3] |
| 1-Methyl-hydantoin | C4H6N2O2 | 114.0429 | [M-H]- | 113.0357 | NA | 113.0352 | 4.42 | 123.64 | NA | |
| 1-Oleoyl-rac-glycerol (MG(18:1(9Z)/0:0/0:0)) | C21H40O4 | 356.2927 | [M+Na]+ | 379.2825 | 197.2 | 379.2815 | 2.64 | 186.04 | 5.66 | [3] |
| 1-palmitoyl-2-myristoyl-sn-glycero-3-phosphocholine (= PC(16:0/14:0) | C38H76NO8P | 705.5309 | [M+Na]+ | 728.5211 | 279.5 | 728.5215 | 0.55 | 291.99 | 4.47 | [4] |
| 1-palmitoyl-2-myristoyl-sn-glycero-3-phosphocholine (= PC(16:0/14:0) | C38H76NO8P | 705.5309 | [M+H]+ | 706.5391 | 278 | 706.5382 | 1.27 | 287.55 | 3.44 | [4] |
| 1-Palmitoyl-2-oleoyl-glycero-3-phosphoethanolamine (16:0/18:1) (= PE( 16:0/18:1) | C39H76NO8P | 717.5309 | [M+Na]+ | 740.5207 | 281.7 | 740.5211 | 0.54 | 291.57 | 3.50 | [3] |
| 1-Palmitoyl-2-oleoyl-glycero-3-phosphoethanolamine (16:0/18:1) (= PE( 16:0/18:1) | C39H76NO8P | 717.5309 | [M+H]+ | 718.5387 | 275.8 | 718.5393 | 0.84 | 283.3 | 2.72 | [3] |
| 1-Palmitoyl-2-stearoyl-glycero-3-phosphocholine (16:0/18:0) | C42H84NO8P | 761.5935 | [M+H]+ | 762.6007 | 290.3 | 762.6014 | 0.92 | 306.39 | 5.54 | [4] |
| 1-Palmitoyl-glycero-3-phosphoethanolamine (16:0) | C21H44NO7P | 453.2855 | [M-H]- | 452.2783 | 209.9 | 452.2782 | 0.22 | 209.98 | 0.04 | [4] |
| 1-Stearoyl-2-palmitoyl-glycero-3-phosphocholine (18:0/16:0) (= PC(18:0/16:0)) | C42H84NO8P | 761.5935 | [M+H]+ | 762.6007 | 290.3 | 762.6014 | 0.92 | 306.39 | 5.54 | [4] |

| | | | | | | | | | | |
|---|---|---|---|---|---|---|---|---|---|---|
| 1,2-Dipalmitoyl-glycero-3-phosphocholine (16:0/16:0) (DPPC) | C40H80NO8P | 733.5622 | [M+H]+ | 734.57 | 284.6 | 734.5701 | 0.14 | 296.15 | 4.06 | [3] |
| 1,2-Dipalmitoyl-glycero-3-phosphocholine (16:0/16:0) (DPPC) | C40H80NO8P | 733.5622 | [M+Na]+ | 756.552 | 286.2 | 756.5526 | 0.79 | 299.81 | 4.76 | [3] |
| 1,2-Distearoyl-sn-glycero-3-phosphate (sodium salt) | C39H76NaO8P | 726.517 | [M+Cl]- | 761.487 | NA | 761.4862 | 1.05 | 318.36 | NA | |
| 1,6-Anhydro-b-glucose | C6H10O5 | 162.0528 | ND | ND | ND | ND | ND | ND | ND | |
| 2-Aminoethylphosphonate | C2H8NO3P | 125.0242 | [M-H]- | 124.0164 | 120.3 | 124.0165 | 0.81 | 125.92 | 4.67 | [3] |
| 2-Deoxyuridine | C9H12N2O5 | 228.0746 | [M+Na]+ | 251.0644 | 150 | 251.0645 | 0.40 | 146.19 | 2.54 | [3] |
| 2-Piperidione | C5H9NO | 99.0684 | [M+H]+ | 100.0757 | NA | 100.0758 | 1.00 | 55.34 | NA | |
| 3-Hydroxy-4-methoxycinnamic acid | C10H10O4 | 194.0579 | [M-H]- | 193.0506 | NA | 193.05 | 3.11 | 138.15 | NA | |
| 3-Hydroxyanthranilic acid | C7H7NO3 | 153.0426 | [M-H]- | 152.0353 | NA | 152.0345 | 5.26 | 128.33 | NA | |
| 3-Phenylpropionate | C9H9O2- | 149.0603 | [M+H]+ | 151.0754 | NA | 151.0762 | 5.30 | 56.69 | NA | |
| 3-Phenylpropionate | C9H9O2- | 149.0603 | [M+Na]+ | 173.0573 | NA | 173.0585 | 6.93 | 147.98 | NA | |
| 3-Ureidopropionate | C4H8N2O3 | 132.0535 | ND | ND | ND | ND | ND | ND | ND | |
| 4-Methylcatechol | C7H8O2 | 124.0524 | [M-H]- | 123.0457 | NA | 123.0451 | 4.88 | 125.38 | NA | |
| 5-Hydroxyindole-3-acetic acid | C10H9NO3 | 191.0582 | [M-H]- | 190.0504 | 137.3 | 190.0505 | 0.53 | 135.53 | 1.29 | [3] |
| Abscisic acid | C15H20O4 | 264.1362 | [M-H]- | 263.1284 | 173.8 | 263.1289 | 1.90 | 161.91 | 6.84 | [3] |
| Adenosine (Adenosyl) | C10H13N5O4 | 267.0968 | [M-H]- | 266.089 | 157.7 | 266.0891 | 0.38 | 151.35 | 4.03 | [3] |
| Adenosine (Adenosyl) | C10H13N5O4 | 267.0968 | [M+H]+ | 268.1046 | 155.5 | 268.1034 | 4.48 | 145.57 | 6.39 | [3] |
| alfa-Linolenic acid | C18H30O2 | 278.2246 | [M-H]- | 277.2168 | 173.9 | 277.2169 | 0.36 | 168.77 | 2.95 | [3] |
| Anandamide C20:4 (AEA) | C22H37NO2 | 347.2824 | [M+H]+ | 348.2897 | 191.63 | 348.2897 | 0.00 | 182.01 | 5.02 | [5] |
| Bilirubin | C33H36N4O6 | 584.2635 | [M-H]- | 583.2562 | 247.18 | 583.2548 | 2.40 | 257.46 | 4.16 | [6] |
| Caffeine | C8H10N4O2 | 194.0804 | [M+H]+ | 195.0861 | 140.2 | 195.0868 | 3.59 | 134.55 | 4.03 | [3] |
| Campesterol | C28H48O | 400.3705 | ND | ND | ND | ND | ND | ND | ND | |
| Chenodeoxycholic acid (CDCA) | C24H40O4 | 392.2927 | [M-H]- | 391.2853 | 209.8 | 391.2845 | 2.04 | 208.22 | 0.75 | [7] |

| | | | | | | | | | | |
|---|---|---|---|---|---|---|---|---|---|---|
| Cholesterol | C27H46O | 386.3549 | [M+H]+ | 387.3631 | 204.6 | 387.3627 | 1.03 | 199.12 | 2.68 | [8] |
| Cholesteryl linoleate | C45H78O2 | 650.6002 | ND | ND | ND | ND | ND | ND | ND | |
| Cholic acid (CA) | C24H40O5 | 408.2876 | [M-H]- | 407.2798 | 203.1 | 407.2798 | 0.00 | 196.35 | 3.32 | [3] |
| Cholic acid (CA) | C24H40O5 | 408.2876 | [M+Na]+ | 431.2774 | 197.7 | 431.2758 | 3.71 | 180.4 | 8.75 | [3] |
| Choline chloride | C5H14ClNO | 139.0764 | [M+Cl]- | 139.0769 | NA | 139.0762 | 5.03 | 133.55 | NA | |
| Cinnamaldehyde | C9H8O | 132.0575 | ND | ND | ND | ND | ND | ND | ND | |
| Coenzyme Q10 | C59H90O4 | 862.6839 | [M-H]- | 861.6755 | NA | 861.6756 | 0.12 | 364.49 | NA | |
| Coenzyme Q10 | C59H90O4 | 862.6839 | [M+H]+ | 863.6912 | NA | 863.6922 | 1.16 | 317.21 | NA | |
| Cytosine | C4H5N3O | 111.0433 | [M+H]+ | 112.0505 | NA | 112.0506 | 0.89 | 54.75 | NA | |
| D-biotin | C10H16N2O3S | 244.0882 | [M+K]+ | 283.0513 | 160.6 | 283.0504 | 3.18 | 147.98 | 7.86 | [3] |
| D-biotin | C10H16N2O3S | 244.0882 | [M+Na]+ | 267.0774 | 161.6 | 267.0754 | 7.49 | 147.49 | 8.73 | [3] |
| Deoxycholic acid (DCA) | C24H40O4 | 392.2927 | [M-H]- | 391.2849 | 202.1 | 391.2845 | 1.02 | 208.22 | 3.03 | [3] |
| Dodecanoic acid (Lauric acid) | C12H24O2 | 200.1776 | [M-H]- | 199.1698 | 154.2 | 199.1694 | 2.01 | 165.16 | 7.11 | [3] |
| Dopamine hydrochloride | C8H11NO2 | 153.079 | [M-H]- | 152.0712 | 133.3 | 152.0715 | 1.97 | 132.72 | 0.44 | [3] |
| Ergocalciferol (vitamine D2) | C28H44O | 396.3392 | [M+H]+ | 397.3465 | 203.2 | 397.3459 | 1.51 | 194.08 | 4.49 | [6] |
| Flavin adenine dinucleotide | C27H33N9O15P2 | 785.1571 | [M-H]- | 784.1493 | 238.5 | 784.1471 | 2.81 | 242.86 | 1.83 | [3] |
| Flavin adenine dinucleotide | C27H33N9O15P2 | 785.1571 | [M-2H+Na]- | 806.1312 | 256.2 | 806.128 | 3.97 | 267.91 | 4.57 | [3] |
| Galactaric acid | C6H10O8 | 210.0376 | [M-2H+Na]- | 231.0117 | 137.5 | 231.0114 | 1.30 | 136.08 | 1.03 | [3] |
| Galactaric acid | C6H10O8 | 210.0376 | [M-H]- | 209.0298 | 132.9 | 209.0295 | 1.44 | 131.74 | 0.87 | [3] |
| Ganglioside GM3 (d18:1/16:0) | C57H104N2O21 | 1152.713 | [M-H]- | 1151.706 | NA | 1151.705 | 0.96 | 138.88 | NA | |
| Ganglioside GM3 (d18:1/18:0) | C52H99N2O24 | 1135.659 | ND | ND | ND | ND | ND | ND | ND | |
| Glyceryl tripalmitate (Tripalmitin) | C51H98O6 | 806.7363 | [M+NH4]+ | 824.7702 | 312.88 | 824.7706 | 0.48 | 336.24 | 7.47 | [5] |
| Glyceryl tripalmitate (Tripalmitin) | C51H98O6 | 806.7363 | [M+Na]+ | 829.7256 | 309.9 | 829.7269 | 1.57 | 331.36 | 6.92 | [5] |
| Glycitein | C16H12O5 | 284.0685 | [M-H]- | 283.0612 | | 283.0608 | 1.41 | 157.59 | | |
| Hesperetin | C16H14O6 | 302.079 | [M-H]- | 301.0712 | 177.9 | 301.0711 | 0.33 | 162.68 | 8.56 | [3] |

| | | | | | | | | | | |
|---|---|---|---|---|---|---|---|---|---|---|
| Hexadecanamide (Palmitamide) | C16H33NO | 255.2562 | [M-H]- | 254.2489 | NA | 254.2492 | 1.18 | 167.94 | NA | |
| Hypotaurine | C2H7NO2S | 109.0197 | ND | ND | ND | ND | ND | ND | ND | |
| Imidazolepropionic acid | C6H8N2O2 | 140.0586 | [M-H]- | 139.0513 | NA | 139.051 | 2.16 | 128.23 | NA | |
| Indole-3-carboxaldehyde | C9H7NO | 145.0528 | [M+H]+ | 146.06 | 125.38 | 146.059 | 6.85 | 128.83 | 2.75 | [9] |
| Inosine | C10H12N4O5 | 268.0808 | [M+Na]+ | 291.071 | 172.1 | 291.0697 | 4.47 | 155.23 | 9.80 | [3] |
| Isovaleric acid (= 3-methylbutanoic acid) | C5H10O2 | 102.0681 | ND | ND | ND | ND | ND | ND | ND | |
| L-Alanine | C3H7NO2 | 89.0477 | ND | ND | ND | ND | ND | ND | ND | |
| L-ascorbic acid | C6H8O6 | 176.0321 | ND | ND | ND | ND | ND | ND | ND | |
| L-Carnosine | C9H14N4O3 | 226.1066 | [M-H]- | 225.0988 | 153.1 | 225.0986 | 0.89 | 154.03 | 0.61 | [3] |
| L-Carnosine | C9H14N4O3 | 226.1066 | [M+Na]+ | 249.0964 | 154.5 | 249.0954 | 4.01 | 144.51 | 6.47 | [3] |
| L-Carnosine | C9H14N4O3 | 226.1066 | [M+H]+ | 227.1144 | 150.8 | 227.1134 | 4.40 | 142.85 | 5.27 | [3] |
| L-gulonic gamma-lactone | C6H10O6 | 178.0477 | [M-H2O-H]- | 159.0299 | 133.7 | 159.0298 | 0.63 | 132.55 | 0.86 | [3] |
| L-gulonic gamma-lactone | C6H10O6 | 178.0477 | [M-H]- | 177.0399 | 129.9 | 177.0394 | 2.82 | 131.49 | 1.22 | [3] |
| L-gulonic gamma-lactone | C6H10O6 | 178.0477 | [M+Na]+ | 201.0375 | 140.9 | 201.0364 | 5.47 | 134.46 | 4.57 | [3] |
| L-Isoleucine | C6H13NO2 | 131.0946 | [M-H]- | 130.0868 | 132.4 | 130.0871 | 2.31 | 132.05 | 0.26 | [3] |
| L-Leucine | C6H13NO2 | 131.0946 | [M-H]- | 130.0868 | 132.5 | 130.0871 | 2.31 | 132.05 | 0.34 | [3] |
| L-Phenylalanine | C9H11NO2 | 165.079 | [M-H]- | 164.0712 | 140.1 | 164.0713 | 0.61 | 137.36 | 1.96 | [3] |
| L-Phenylalanine | C9H11NO2 | 165.079 | [M-H]- | 164.0717 | 140.6 | 164.0713 | 2.44 | 137.36 | 2.30 | [3] |
| L-Threonine | C4H9NO3 | 119.0582 | ND | ND | ND | ND | ND | ND | ND | |
| L-Tyrosine | C9H11NO3 | 181.0739 | [M-H]- | 180.0666 | 145 | 180.0662 | 2.22 | 139.68 | 3.67 | [3] |
| Linoleic acid | C18H32O2 | 280.2402 | [M-H]- | 279.2324 | 174.6 | 279.2317 | 2.51 | 169.32 | 3.02 | [3] |
| Lithocholic acid (LCA) | C24H40O3 | 376.2977 | [M-H]- | 375.2899 | 201.4 | 375.2895 | 1.07 | 201.12 | 0.14 | [3] |
| Malic acid | C4H6O5 | 134.0215 | ND | ND | ND | ND | ND | ND | ND | |
| Margaric acid | C17H34O2 | 270.2559 | [M-H]- | 269.2486 | 173.1 | 269.248 | 2.23 | 176.6 | 2.02 | [3] |
| Myristic acid | C14H28O2 | 228.2089 | [M-H]- | 227.2016 | NA | 227.2018 | 0.88 | 158.89 | NA | |
| N-Acetylagmatine (acetate salt) | C7H16N4O | 172.1324 | [M+Cl]- | 207.1018 | NA | 207.1014 | 1.93 | 146.07 | NA | |

| | | | | | | | | | | |
|---|---|---|---|---|---|---|---|---|---|---|
| N-Acetylagmatine (acetate salt) | C7H16N4O | 172.1324 | [M+H]+ | 173.1397 | NA | 173.1413 | 9.24 | 58.79 | NA | |
| N-Acetylputrescine | C6H14N2O | 130.1106 | [M+H]+ | 131.1184 | NA | 131.1196 | 9.15 | 88.64 | NA | |
| N-oleoyl-D-erythro-sphingosine (Cer d18:1/18:1) | C42H79NO8 | 725.5806 | [M+Cl]- | 598.4971 | NA | 598.4965 | 1.00 | 271.11 | NA | |
| N-oleoyl-D-erythro-sphingosine (Cer d18:1/18:1) | C42H79NO8 | 725.5806 | [M-H]- | 562.5205 | NA | 562.5192 | 2.31 | 268.22 | NA | |
| N-palmitoyl-D-erythro-sphingosine (Cer d18:1/16:0) | C34H67NO3 | 537.5121 | [M-H]- | 536.5048 | 244.6 | 536.5037 | 2.05 | 264.5 | 8.14 | [4] |
| Nicotinic acid | C6H5NO2 | 123.032 | [M+H]+ | 124.0398 | 123.9 | 124.04 | 1.61 | 125.09 | 0.96 | [3] |
| Oleic acid | C18H34O2 | 282.2559 | [M+Na]+ | 305.2457 | 180.2 | 305.2472 | 4.91 | 169.66 | 5.85 | [3] |
| Oleoyl-L-carnitine | C25H47NO4 | 425.3505 | [M+Na]+ | 448.3403 | 227.2 | 448.34 | 0.67 | 211.99 | 6.69 | [3] |
| Oleoyl-L-carnitine | C25H47NO4 | 425.3505 | [M+H]+ | 426.3583 | 217.3 | 426.3575 | 1.88 | 207.94 | 4.31 | [3] |
| Palmitic acid (= hexadecanoic acid) | C16H32O2 | 256.2402 | [M-H]- | 255.2329 | 169 | 255.2329 | 0.00 | 165.53 | 2.05 | [3] |
| Palmitoyl-L-carnitine | C23H45NO4 | 399.3349 | [M+Na]+ | 422.3247 | 228.9 | 422.3243 | 0.95 | 212.6 | 7.12 | [3] |
| Palmitoyl-L-carnitine | C23H45NO4 | 399.3349 | [M+H]+ | 400.3427 | 214.7 | 400.3423 | 1.00 | 204.79 | 4.62 | [3] |
| Pentadecanoic acid | C15H30O2 | 242.2246 | [M-H]- | 241.2168 | 165.1 | 241.2173 | 2.07 | 177.16 | 7.30 | [3] |
| Piperine | C17H19NO3 | 285.1365 | [M+H]+ | 286.1438 | 171.4 | 286.1439 | 0.35 | 160.99 | 6.07 | [8] |
| Piperine | C17H19NO3 | 285.1365 | [M+Na]+ | 308.1263 | 166.9 | 308.1253 | 3.25 | 173.17 | 3.76 | [9] |
| Piperonyl acetone | C11H10O3 | 190.063 | [M-H]- | 191.0713 | NA | 191.0713 | 0.00 | 140.55 | NA | |
| Pregnenolone sulfate sodium | C21H31NaO5S | 418.179 | [M+Cl]- | 453.1484 | NA | 453.1468 | 3.53 | 219.96 | NA | |
| Prostaglandine E2 | C20H32O5 | 352.225 | [M-H]- | 351.2177 | 192.06 | 351.2172 | 1.42 | 195.87 | 1.98 | [10] |
| Prostaglandine E2 | C20H32O5 | 352.225 | [M+Na]+ | 375.2142 | 196.8 | 375.2163 | 5.60 | 182.4 | 7.32 | [6] |
| Prostaglandine E2 | C20H32O5 | 352.225 | [M-H2O-H]- | 333.2071 | 183.57 | 333.209 | 5.70 | 181.6 | 1.07 | [6] |
| Pseudouridine | C9H12N2O6 | 244.0695 | [M-H]- | 243.0617 | 150.3 | 243.0617 | 0.00 | 151.65 | 0.90 | [3] |
| Pyrrole-2-carboxylic acid | C5H5NO2 | 111.032 | ND | NA | NA | NA | NA | NA | NA | |
| Pyruvic acid | C3H4O3 | 88.016 | ND | NA | NA | NA | NA | NA | NA | |
| Retinol acetate | C22H32O2 | 328.2402 | ND | NA | NA | NA | NA | NA | NA | |

| | | | | | | | | | | |
|---|---|---|---|---|---|---|---|---|---|---|
| S-(5'-Adenosyl)-L-methionine | C15H22N6O5S | 398.1372 | [M+H]+ | 399.145 | 185.6 | 399.1443 | 1.75 | 170.98 | 7.88 | [3] |
| Saccharin | C7H5NO3S | 182.999 | [M-H]- | 181.9917 | NA | 181.9915 | 1.10 | 129.76 | NA | |
| Sodium glycochenodeoxycholate (GCDCA) | C26H43NO5 | 449.3141 | [M-H]- | 448.3063 | 200.6 | 448.3068 | 1.12 | 192.82 | 3.88 | [3] |
| Sodium glycodeoxycholate (GDCA) | C26H43NO5 | 449.3141 | [M+Cl]- | 506.2655 | NA | 506.2646 | 1.78 | 207.24 | NA | |
| Spermidine | C7H19N3 | 145.1579 | [M+H]+ | 146.1652 | NA | 146.1656 | 2.74 | 104.99 | NA | |
| Stearic acid | C18H36O2 | 284.2715 | [M-H]- | 283.2642 | 177.3 | 283.2637 | 1.77 | 173.56 | 2.11 | [3] |
| Taurine | C2H7NO3S | 125.0147 | [M-H]- | 124.0074 | **NA** | 124.0074 | 0.00 | 122.66 | NA | |
| Theobromine | C7H8N4O2 | 180.0647 | [M+H]+ | 181.0721 | 131.1 | 181.0709 | 6.63 | 131.96 | 0.66 | [3] |
| Theophylline | C7H8N4O2 | 180.0647 | [M+H]+ | 181.0709 | 140.9 | 181.0709 | 0.00 | 131.96 | 6.34 | [3] |
| Trehalose | C12H22O11 | 342.1162 | [M-H]- | 341.1084 | 169.9 | 341.1074 | 2.93 | 163.49 | 3.77 | [3] |
| Trehalose | C12H22O11 | 342.1162 | [M+Na]+ | 365.106 | 176.1 | 365.1048 | 3.29 | 165.25 | 6.16 | [3] |
| Tridecanoic acid | C13H26O2 | 214.1933 | [M-H]- | 213.186 | 157.9 | 213.1853 | 3.28 | 156.88 | 0.65 | [4] |
| Tryptamine | C10H12N2 | 160.1 | [M-H]- | 159.0928 | NA | 159.092 | 5.03 | 134.71 | NA | |
| Valeric acid | C5H10O2 | 102.0681 | ND | ND | ND | ND | ND | ND | ND | |
| Xanthine | C5H4N4O2 | 152.0334 | [M-H]- | 151.0261 | NA | 151.0262 | 0.66 | 123.2 | NA | |
| Xanthurenic acid | C10H7NO4 | 205.0375 | [M+H]+ | 206.0453 | 140.4 | 206.0438 | 7.28 | 134.82 | 3.97 | [3] |

**ND - not detected**
**NA - not available**
